# Genome-wide family prediction unveils molecular mechanisms underlying the regulation of agronomic traits in *Urochloa ruziziensis*

**DOI:** 10.1101/2023.09.25.559305

**Authors:** Felipe Bitencourt Martins, Alexandre Hild Aono, Aline da Costa Lima Moraes, Rebecca Caroline Ulbricht Ferreira, Mariane de Mendonça Vilela, Marco Pessoa-Filho, Mariana Rodrigues Motta, Rosangela Maria Simeão, Anete Pereira de Souza

## Abstract

Tropical forage grasses, especially species of the genus *Urochloa*, play an important role in cattle production and are the main food source for animals in tropical/subtropical regions. Most of the species are apomictic and tetraploid, which gives special importance to *U. ruziziensis*, a sexual diploid species that can be tetraploidized for use in interspecific crosses with apomictic species. As a means to assist in breeding programs, this study investigates the applicability of genome-wide family prediction (GWFP) in *U. ruziziensis* half-sibling families to predict growth and biomass production. Machine learning and feature selection algorithms were used to reduce the necessary number of markers for prediction and to enhance the predictive ability across the phenotypes. Beyond that, to investigate the regulation of agronomic traits, the positions of the markers with more importance for the prediction were considered putatively associated to quantitative trait loci (QTLs), and in a multiomic approach, genes obtained in the species transcriptome were mapped and linked to those markers. Furthermore, a gene coexpression network was modeled, enabling the investigation of not only the mapped genes but also their coexpressed genes. The functional annotation showed that the mapped genes are mainly associated with auxin transport and biosynthesis of lignin, flavonol and folic acid, while the coexpressed genes are associated with DNA metabolism, stress response and circadian rhythm. The results provide a viable marker-assisted breeding approach for tropical forages and identify target regions for future molecular studies on these agronomic traits.

## 1 Introduction

Pastures composed of tropical forage grasses, particularly those belonging to the *Urochloa* genus, serve as the main food source for livestock animals in tropical and subtropical regions. These pastures play a significant role in the economic sectors associated with beef and dairy production, as well as seed markets (Jank et al., 2014; Ferreira et al, 2021). The genetic improvement of *Urochloa* species is recent, starting approximately 40 years ago, and presents challenges due to varying ploidy levels, high heterozygosity, and a prevalent mode of reproduction through apomixis (Ferreira et al., 2021; Simeão et al., 2021). Among the main goals of breeding programs are the development of cultivars that exhibit tolerance to biotic stresses, adaptability to future climate changes, and increased productivity with enhanced nutritional value to optimize animal performance (Pereira et al., 2018b; Simeão et al., 2021).

These goals can be expedited through the incorporation of genomic selection (GS) into breeding cycles. GS employs statistical models to perform genomic predictions (GPs) of plant performance based on genetic markers, mainly single nucleotide polymorphisms (SNPs) (Daetwyler et al, 2013). Although the estimation of GP models has already demonstrated feasibility in other important polyploid crops (de Bem Oliveira et al., 2020; Pincot et al., 2020; Ferrão et al., 2021; Haile et al., 2021; Juliana et al., 2022; Petrasch et al., 2022), this methodology has only recently started to be tested in *Urochloa* spp. (Matias et al., 2019a; Aono et al., 2022). Therefore, efforts must be directed toward the establishment of high-quality marker panels and large-scale phenotyping (Simeão et al., 2021). Fortunately, two *Urochloa* spp. genomes, specifically *U. ruziziensis* (2n=2x=18), have recently become available (Pessoa-Filho et al., 2019; Worthington et al., 2021), facilitating the identification of many SNPs with the potential to enhance the accuracy of GP analyses in *Urochloa* spp.

Traditionally, GP models employ a dense dataset of molecular markers to compute genomic estimated breeding values at the individual level (Meuwissen et al., 2001). However, in the case of *U. ruziziensis* and other forage species, such as alfalfa and ryegrass, it is a common practice to employ the family (full or half-siblings) as the basic unit for phenotyping and selection (Simeão et al., 2012; Simeão et al., 2016a; Simeão et al., 2016b; Biazzi et al., 2017; Cericola et al., 2018; Jia et al., 2018; Andrade et al., 2022). This practice makes the development of genome-wide family prediction (GWFP) approaches highly advantageous. By considering family groups as the measurement unit, there is a reduction in genotyping efforts, as well as the costs associated with developing GP models (Zou et al., 2016; Rios et al., 2021; Murad Leite Andrade et al., 2022). Furthermore, the implementation of GWFP can improve the predictive ability of selection, increasing the rate of genetic gains for complex traits, as demonstrated in studies on loblolly pine and alfalfa (Rios et al., 2021; Murad Leite Andrade et al., 2022).

To identify family-pool markers, sequencing approaches can be employed to generate a large number of SNP markers (Elshire et al., 2011; Poland et al., 2012). Genotyping-by-sequencing (GBS) is a cost-effective and high-throughput genotyping method that can be used to identify SNPs even in the absence of a reference genome. However, it is important to ensure a reasonable sequencing depth to minimize the occurrence of missing data points (Thakral et al., 2022). GBS has been employed in several studies on family-pool genotyping (Futschik & Schlötterer, 2010; Bélanger et al., 2016; Cericola et al., 2018; Schneider et al., 2022) due to its advantages and straightforward applicability in obtaining allele counts from sequencing reads (Byrne et al., 2013). Consequently, in the context of family-pool GP, the use of allele counts derived from GBS allows for direct inference without the need for estimating allelic dosages (Guo et al., 2018).

In addition to the application of GP models in GS approaches, family-pool markers can also be employed in genome-wide association studies (GWAS). Unlike selection-based applications, GWAS aims to identify loci that are associated with a greater extent of genetic variation, thereby enhancing the understanding of the genetic architecture underlying complex traits (Ashraf et al., 2014; Zhang et al., 2014; Fé et al. 2015). In this sense, adopting a family-based approach provides a more comprehensive perspective on the genetic variations related to the configuration of traits across different families. Once these genomic associations have been assessed, additional omics approaches can be employed to further elucidate the biological mechanisms triggered by adjacent genes and their association with the configuration of complex traits (Scossa et al., 2021).

Traditionally, data generated from various levels of biological information, such as genomics, transcriptomics, and proteomics, have been analyzed separately. However, more recently, the integration of data followed by appropriate statistical analysis has emerged as a promising approach to unravel the biological implications of different traits in humans (Yang et al, 2014), microorganisms (Borin et al., 2018; Rosolen et al., 2022), animals (Parker Gaddis et al., 2016; Mateescu et al., 2017), and plants (Francisco et al., 2021; Cardoso-Silva et al., 2022). Despite the economic importance of *U. ruziziensis* and the availability of molecular data resources, no study incorporating multiomics has been conducted on *U. ruziziensis* or any species of the *Urochloa* genus.

Although assessing different aspects, GP and GWAS possess complementary advantages, providing robust information for the identification of potential candidate genes related to agronomically important traits. Methodologies originally used for GP have been applied in GWAS to detect loci associated with the trait of interest (Goddard et al., 2016; Wang et al., 2020; Wolc et al., 2022). Conversely, association studies have demonstrated their usefulness in enhancing GP (Zhang et al., 2014; Bian et al., 2017; Jeong et al., 2020). To further enhance the outcomes of association and prediction studies, researchers have explored the integration of machine learning (ML) algorithms. Despite the controversial incorporation of ML in GP, with some studies highlighting its advantages (Ma et al., 2018; Waldman et al., 2020; Aono et al., 2022) and others refuting them (Montesinos-López et al., 2019; Zingaretti et al., 2020; Crossa et al., 2019), numerous investigations consistently demonstrate that ML-based strategies incorporating feature selection (FS) techniques effectively reduce marker density. These methods not only maintain or enhance prediction accuracy but also enable the identification of polymorphisms associated with phenotypes (Li et al., 2018; Aono et al., 2020; Pimenta et al., 2021; Aono et al., 2022).

In this study, we assessed the feasibility of family-based genotyping in autotetraploid *U. ruziziensis* (2n = 4x = 36) and investigated the GWFP capability to predict biomass production and growth traits in both wet and dry seasons. We employed traditional statistical methods as well as ML algorithms to analyze the data. To enhance prediction accuracy, we employed FS strategies to identify subsets of SNP markers with increased predictive power. Furthermore, we used an ML tree-based approach to estimate the importance of these variations in prediction. The most significant markers were then used as a guide to map RNA-Seq assembled genes, which were considered putatively associated with the investigated traits. To gain a deeper understanding of the molecular mechanisms underlying the regulation of these traits in the different seasons investigated, we expanded the set of identified genes by constructing a gene coexpression network (GCN). Our study not only brings innovation to GWFP, but also proposes a means of integrating genomic and transcriptomic data. Moreover, our findings contribute to the expansion of knowledge on the biological processes influencing the investigated agronomic traits. The outcomes of this work offer valuable resources for future studies and breeding programs targeting the *Urochloa* genus.

## 2 Materials and Methods

### 2.1 Urochloa ruziziensis phenotyping

The progenies used in this study were generated as part of the *Urochloa* breeding program of the Brazilian Agricultural Research Corporation (Embrapa) Beef Cattle (EBC), located in Campo Grande, Mato Grosso do Sul State, Brazil (20°27’S, 54°37’W, 530 m), as described by Simeão et al. (2012, 2016a, b). In 2010, seven sexual autotetraploid-induced accessions (R30, R38, R41, R44, R46, R47 and R50) were replicated 20 times to create an open pollination randomized field organized into 26 lines and 12 columns spaced by 2 meters. In 2012, out of the 140 plants, 59 were selected to form breeding progenies and compose the experiment of the study. This selection was based on their viable seed production and flowering synchrony. A total of 1,180 individuals (20 seeds from each of the 59 plants selected) were planted in a randomized block design, with one plant per plot spaced 1.5 m apart (Simeão et al., 2016a, b). From the 59 half-sibling progenies, 50 were chosen based on the criterion of selecting the progenies with more plants that succeeded in the field.

The phenotypic evaluations were performed considering nine clippings at 15 cm height: (1) March 2012; (2) January 2013; (3) April 2013; (4) May 2013; (5) September 2013; (6) October 2013; (7) November 2013; (8) December 2013; and (9) January 2014. According to the climatological water balance assessed through the available water capacity (AWC) metric (Supplementary Fig. S1) (Simeao et al, 2016a, b), six clippings were performed in the wet season (1-3,7-9) and three in the dry season (4-6). In addition to the nine clippings, we had a total sum (T) evaluation for each phenotype in the period.

The agronomic traits evaluated in all clippings were green matter yield (GM) and dry matter yield (DM), both measured in grams per family, and regrowth (RG), with scores varying from 0 to 6 as described by Figueiredo et al. (2012). In addition, in clippings 2 and 5, approximately 200 g of leaves and stems from each plant were used to estimate leaf dry matter yield (LDM) and stem dry matter yield (SDM). Considering that clipping 1 was discarded from the analysis, we evaluated 33 combinations of agronomic traits and clippings (clippings 2-9 for GM, DM and RG, and clippings 2 and 5 also for SDM and LDM) (Fig. 1.1), which we considered different phenotypes.

**Figure 1.**
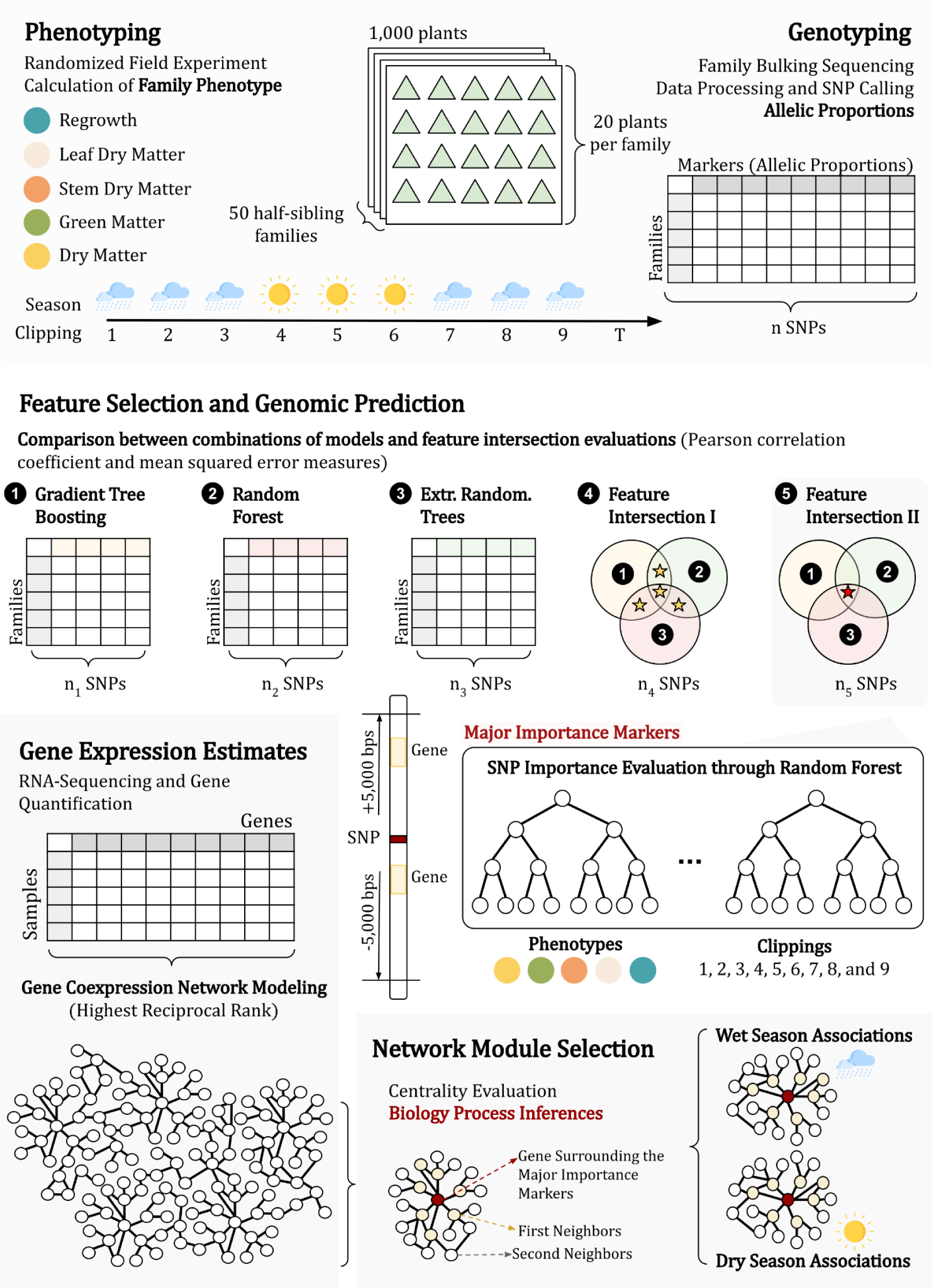
The approach established in this research can be divided into three main parts: (i) phenotyping and genotyping the population (1); (ii) identifying phenotypically associated markers through genomic prediction (2 and 3); and (iii) investigating the genes physically linked to the markers in a coexpression network (4, 5 and 6).

For each combination of agronomic traits and clippings, we employed the following linear mixed-effects model:

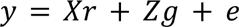

where *y* represents the phenotypic measurements, X is the design matrix of the fixed repetition effects *r*, Z is the design matrix of the random genotypic effects *g*, and *e* is the random residual vector. All the statistical analyses were performed using the software Selegen - REML/BLUP (Resende 2002; Colombari-Filho et al. 2013). Narrow-sense trait heritability estimates were corrected using the Wright’s coefficient of relationship, as described by Simeão et al. (2016a).

To obtain family measurements, we calculated the average of each trait per family and scaled the results between 0 and 1 with the Min-Max technique. To perform a data descriptive analysis of family traits, we used boxplots to assess the distribution and outliers, computed Pearson’s correlation among all phenotype clippings and performed a principal component analysis (PCA) to assess population structure. The descriptive analysis was performed in R (R Core Team, 2021), and all PCA and plots were performed with the package pcaMethods (Stacklies et al., 2007) and the package ggplot2 (Wickham & Chang, 2016), respectively.

### 2.2 Genotyping

Genomic DNA of all individuals was extracted using the DNeasy Plant kit (QIAGEN) and pooled according to each family, totaling 50 samples. GBS libraries were constructed following the method proposed by Poland et al. (2012) using a combination of a rare cutting enzyme (EcoT22I) and a frequent cutting enzyme (MspI). Subsequently, libraries were sequenced as 150-bp single-end reads using the High Output v2 Kit (Illumina, San Diego, CA, USA) in a NextSeq 500 platform (Illumina, San Diego, CA, USA).

We performed quality evaluation of GBS raw sequence reads using FastQC version 0.11.5 (Andrews, S. 2010) and SNP calling using the TASSEL-GBS pipeline (Glaubitz et al., 2014) modified for polyploids (Pereira et al., 2018a). The reads were aligned to the *U. ruziziensis* genome assembly (Pessoa-Filho et al, 2019; GenBank Assembly GCA_015476505.1) using the BowTie 2.3.1 aligner (Langmead et al., 2009), and only uniquely mapped reads were employed. SNP markers were filtered using VCFtools v0.1.17 (Danecek et al., 2011) with the following criteria: a minimum sequencing depth of 20 reads, no more than 25% missing data per site, biallelic SNPs only, and removal of redundant (same genotypes in all samples) markers from the sets (Fig. 1.1).

The allele frequency for each marker was estimated as the ratio between the number of reads for the alternative allele and the total number of reads. Missing data were replaced by the site mean of allele frequency. Furthermore, a PCA was performed on the complete genotype data to assess population structure.

### 2.3 Genomic Prediction and Feature Selection

To create subsets of markers for each phenotype, three FS techniques were applied to the SNP data using the Python 3 library scikit-learn v1.0.2 (Pedregosa et al., 2011): gradient tree boosting (FS-1) (Chen & Guestrin, 2016), extremely randomized trees (FS-2) (Geurts, Ernst & Wehenkel, 2008), and random forest (FS-3) (Breiman, 2001). For the FS-1 technique, we employed the mean squared error (MSE) as the loss function, set the learning rate to 0.1, and considered 100 boosting stages. The criterion for assessing split quality was based on the MSE with improvement score by Friedman. We established that a minimum of 2 samples was required to split an internal node, while a minimum number of 1 sample was required for a leaf node. Furthermore, we constrained the maximum number of nodes within the trees to 3. For FS-2 and FS-3, the forest consisted of 100 trees, employing the MSE as the quality measurement function. The minimum number of samples required to split an internal node and the minimum number of samples required to form a leaf node were consistent with those of FS-1. In FS-3, the trees had no node limit, and bootstrapping was employed. Then, to perform modeling, we created feature intersection (FI) datasets by evaluating the intersection of the FS methods, considering markers that were selected by at least two FS techniques (FI-1) and markers that were selected by all three FS techniques (FI-2), similar to the approach proposed by Aono et al. (2020) and Aono et al. (2022) (Fig. 1.2).

As GP strategies, we estimated different models considering six regression approaches across the 33 combinations of traits and clippings, as well as both the reduced (FI-1 and FI-2) and complete versions of the dataset. As conventional GP models, we employed the semiparametric reproducing kernel Hilbert space (RKHS) with a Gaussian kernel (GK) as the covariance function using the R package BGGE v0.6.5 (Granato et al., 2018) and Bayesian ridge regression (BRR) with the R package BGLR v1.0.9 (Perez & de los Campos, 2014). Both models were estimated using 20,000 iterations with a thinning of 5 and a burn-in of 2,000. Additionally, we evaluated four ML algorithms using Python 3 with the scikit-learn library v1.0.2 (Pedregosa et al., 2011): (i) support vector machine (SVM) (Cristianini & Shawe-Taylor, 2000); (ii) random forest (RF) (Breiman, 2001); (iii) adaptive boosting (AB) (Freund & Schapire, 1997); and (iv) multilayer perceptron (MLP) neural network (Popescu et al., 2009). For SVM regression, a radial basis function was used as the kernel, with the gamma coefficient defined as 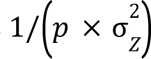, where *p* represents the number of loci and 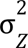 the variance of the genotype matrix **Z**. The RF regression was performed with the same parameters as those described for FS-3. For AB, we employed a decision tree regressor as the base estimator, used a linear loss function to assign weights, and limited the maximum boosting interaction to 50 estimators. Finally, the MLP neural network was constructed with a single hidden layer comprising 100 neurons activated by the rectified linear unit (ReLU) function. We employed a quasi-Newton method to optimize the weights and applied a regularization term of 0.001 strength in the L2 regularization term.

The evaluation of the previously described models for GP was performed using a k-fold (k=5) cross validation strategy, repeated 100 times. Two metrics were measured: predictive ability (PA), quantified as the Pearson correlation coefficient, and MSE (Fig. 1.3).

To compare the models, the phenotype clippings and the datasets, we used ANOVAs with multiple comparisons by Tukey’s tests implemented in the agricolae R package (De Mendiburu & De Mendiburu, 2020) (Fig. 1.3). For PA, we considered the best scenario to be that in which Tukey’s test had “a” or “a” combined with other letters, such as “ab” or “abc”, which represents the highest values. On the other hand, for MSE, a scenario is better when its MSE value is lower. Therefore, we considered the best scenarios those with the higher letter or combined with other letters (i.e., “f”, “ef” or “def”).

### 2.4 Major importance markers

After identifying the best dataset of markers for each phenotype clipping, we used the random forest algorithm (Breiman, 2001) to estimate the impurity importance of each SNP marker. This estimation was performed considering the Gini importance, which quantifies the normalized total reduction in the criterion (MSE) achieved by each feature (the sum of the feature importance across all markers is equal to 1). To obtain a more refined subset comprising only the markers most likely associated with agronomic traits, we established the major importance set by selecting the top 3 Gini importance markers in each phenotype clipping (Fig. 1.3). In cases where the sum of these three values did not reach 0.5, we continued selecting additional values until the condition was satisfied. Furthermore, a PCA was performed using the major importance markers dataset.

### 2.5 Transcriptome assembly, quantification and annotation

Previous RNA-Seq data of 11 genotypes of *U. ruziziensis* were used to assess gene expression (Hanley et al., 2021; NCBI BioProject PRJNA513453). Raw data were quality-trimmed using Trimmomatic v0.39 (Bolger, Lohse & Usadel, 2014). The Illumina adapters, the first 12 bases of the read, and the leading and trailing bases with quality less than 3 were trimmed; the sliding window of 4 bases was set to cut the read when quality/base was less than 20 and only reads with more than 75 bases were kept. Then, the filtered reads were de novo assembled by Trinity v2.5.1 (Grabherr et al., 2011) considering a minimum contig length of 300 bases, and assembly integrity was evaluated using the Trinity.pl package utility (Fig. 1.4).

SALMON 1.1.0 (Patro et al., 2017) was used to quantify transcript expression, which was subsequently summarized at the gene level using the tximport R package (Soneson et al., 2015). We retained only genes with more than one transcript per million (TPM) in at least three of the 11 samples, disregarding genes with low-level expression. The longest isoform for each gene was selected, and BUSCO v5.2.2 (Manni et al., 2021) was used to evaluate the annotation completeness against the Viridiplantae database. Finally, we aligned the filtered assembly to the UniProt database (Bateman et al., 2020) using Blastx and Blastn 2.10.0 (Altschul et al., 1990) with an e-value cutoff of 1e-10. Gene Ontology (GO) terms were retrieved using Trinotate software (Bryant et al., 2017), which performed functional annotation (Fig. 1.4).

### 2.6 Genes linked with markers and GO enrichment

To identify genes physically linked to major importance markers (section 2.4), we conducted alignments between the genes derived from the transcriptome assembly (section 2.5) against the *U. ruziziensis* genome (Pessoa-Filho et al., 2019). Therefore, genes that aligned in a window of 5,000 bp up- and downstream of the marker position were considered physically linked. The alignment was performed using Blastn 2.10.0 (Altschul et al., 1990) with a minimum query coverage of 75% and an E-value cutoff of 1e-6. To visualize the gene position within the genome, we constructed a physical map using MapChart v2.32 (Voorrips, 2002), including information regarding the phenotype and the seasonal associations, as well as Gini importance (Fig. 1.5). In addition, a circular map was constructed using the R package circlize v0.4.14 (Gu et al., 2014) to show the associated genes that were duplicated.

Finally, to obtain a functional profile of the genes linked to the markers, biological process GO term enrichment analysis was performed. This step was achieved with the R package topGO (Alexa & Rahnenfuhrer, 2022), and GO terms with p values < 0.01 in Fisher’s exact test were considered significantly enriched.

### 2.7 Coexpression network

We modeled a GCN using the transcript quantifications normalized in transcripts per million (TPM) and the highest reciprocal rank (HRR) (Mutwil et al., 2010) approach, considering a limit of 30 edges. From the GCN, we selected the genes associated with the agronomic traits and included highly correlated genes that were not considered in the network ranking (Pearson correlation coefficient ≥ 0.9 and a maximum p value of 0.01 with Bonferroni correction). From this defined gene set, we selected the first and second gene neighbors in the GCN. To evidence the gene associations with the two seasons, we highlighted genes related to phenotype clippings 2,3,7,8 and 9, considering them as components of a wet-season associated network. Similarly, genes associated with clippings 4, 5 and 6 were selected to form the dry-season associated network.

Network visualization and evaluation were performed using Cytoscape software v3.9.1 (Shannon et al., 2003). For each gene, we calculated the degree centrality measure with the methods of Barabási & Oltvai, 2004, and considered the genes with outlier values as hubs. Finally, biological process GO term enrichment analyses were performed for the selected genes, including first and second neighbors, to produce a general and seasonal functional profile of the metabolic pathways associated with the agronomic traits with the same method described in 2.6 (Fig. 1.6).

## 3 Results

### 3.1 Phenotypic and genotypic data analyses

In our study, we evaluated five important traits for forage grasses (GM, DM, RG, LDM, and STM) across various clippings selected based on wet and dry seasons. Individual measurements were averaged at the subfamily level, and we excluded data from the first clipping. The descriptive analysis of subfamily based phenotypic data did not reveal any discernible patterns concerning the dispersion and skewness of the traits (Supplementary Fig. S2). We did not identify any outliers in 17 out of the 33 traits. Despite the absence of any apparent similarity in phenotypic dispersion between the phenotypes evaluated, the correlation analysis yielded significant values for all the comparisons conducted (Supplementary Fig. S3). We observed an average R Pearson correlation coefficient of 0.72 (Supplementary Fig. S3), with the strongest correlations (∼1) observed between the same clippings of GM and DM. Additionally, early clippings (2 and 3) tended to be less correlated with all other measures. This pattern was particularly more pronounced for GM, DM, and SDM. In contrast, SDM in clipping 2 exhibited the lowest correlation with all other phenotypes (Supplementary Fig. S3). The progeny mean narrow-sense heritabilities for all phenotype clippings showed a mean value of 0.79, ranging from 0.44 (SDM in clipping 2) to 0.92 (LDM in clipping 5) (Supplementary Table S1).

The GBS experiment generated ∼720 million reads, which were processed into 1.3 million tags using the Tassel pipeline. We identified a total of 77,413 SNP markers in this step. After applying quality filters, estimating allele frequencies, and imputing missing genotypes, we retained 28,106 of these markers. This final dataset of markers is referred to as the “complete data” (CD).

By using the phenotypic and genotypic data, we performed PCAs, plotting the dispersion of subfamilies using the scores of the first two principal components (PCs) (Supplementary Figs. S4-S5). Although arising from different sources of variation (the proportion of variance explained by the first two PCs was 85.2% and 57.2% for the phenotypic and genotypic data, respectively), similar patterns could be observed. To corroborate such a similarity, we colored the samples from the genotypic PCA scatter plot using PC1 of the phenotypic data. Even without a pronounced presence of 3 groups, as in the phenotypic PCA, the coloring in the genotypic PCA evidenced a clear association between both PCA results (Supplementary Fig. S5).

### 3.2 Genome-wide family prediction

The predictive performance of the GP models at the family level using the CD was assessed through the consideration of two conventional approaches (RKHS and BRR) across 33 phenotypes. Employing a 100-times 5-fold CV strategy, the RKHS model exhibited slightly superior results compared to BRR, with a mean PA of ∼0.762 and mean MSE of ∼0.025, contrasted to a mean PA of ∼0.745 and a mean MSE of 0.026 in BRR. We observed a maximum PA of ∼0.875 in the DM-8 trait and a minimum PA of ∼0.490 in SDM-2. Aiming to achieve higher performance levels, we evaluated four ML algorithms (SVM, RF, AB and MLP). Among these models, SVM exhibited the best overall performance, with a mean PA of ∼0.759 and a mean MSE of ∼0.026; however, it did not surpass the performance of the RKHS approach. By considering Tukey’s test results for MSE, it became evident that the RKHS model significantly outperformed SVM, emerging as the superior approach in 30 traits compared to 13 of SVM (Supplementary Table S2-4). Our results indicate that when using CD for prediction, the ML algorithms were unable to outperform the performance of conventional models.

To increase our predictive accuracy and assess potential associations between traits and markers, we selected specific subsets of SNPs for each of the 33 traits based on the intersections established between FS sets. Each FS approach yielded a distinct quantity of markers: FS-1 selected sets with quantities ranging from 129 to 175 markers (mean of ∼150, 0.53% of the CD); FS-2 from 484 to 1154 (mean of ∼848, 3% of the CD); and FS-3 from 563 to 853 (mean of ∼699, 2.5% of the CD). By considering the intersection approaches established, we obtained FI-1 with SNP quantities ranging from 76 to 122 markers (mean of ∼102, 0.36% of the CD) and FI-2 with quantities varying from 5 to 23 markers (mean ∼11, 0.04% of the CD) (Supplementary Table S5). In addition to obtaining more restricted sets, these markers selected by FI have more evidence of trait associations, as they were selected by multiple algorithms. In this sense, model performances using the CD were contrasted with the use of models created from the datasets selected by FI-1 and FI-2.

The employment of the FI datasets increased the performance of all models for all traits. This improvement was particularly pronounced in the AB and RF models, which presented the highest levels of accuracy, overcoming RKHS in both FI sets. Among the six models evaluated, the FI-1 approach presented an improved overall performance when compared to FI-2, being considered by Tukey’s test the best approach in 168 (FI-2 = 100) and 136 (FI-2 = 89) scenarios for PA and MSE, respectively (Supplementary Table S6). However, individual results for the best models in each scenario were similar, as indicated by the best model in FI-1 (AB with a mean PA of ∼0.894 and a mean MSE of ∼0.013) and FI-2 (RF with a mean PA of ∼0.893 and a mean MSE of ∼0.013) (Supplementary Table S2-4). Furthermore, when analyzing the clippings of a phenotype, we observed that the best performances for clippings in the combinations AB-FI-1 and RF-FI-2 varied in GM and DM, but for RG (clipping 3), SDM (clipping 5) and LDM (clipping 5), the results were equivalent (Supplementary Tables S2-3 and 7).

In this sense, we observed that for the prediction task, both combinations AB-FI-1 and RF-FI-2 can be employed with comparable performance levels. However, for investigating trait‒marker associations and catalogs of putative associated genes, FI-2 represents a more restrictive approach. With sets (mean of ∼11 markers) approximately ten times smaller than the sets of FI-1 (mean of ∼102 markers), FI-2 markers provide a group of markers with a probable reduced number of false positive associations. Therefore, we considered the combination RF-FI-2 as the most promising approach to be employed in our datasets. In addition to the significant decrease in marker density through FI-2, the RF algorithm demonstrated high efficiency for prediction with a PA increase of 6.9% and an MSE reduction of 22.6% when compared to the RKHS using the FI-2 dataset or 17% when compared to the RKHS using the CD dataset (Table 1).

**Table 1.** Comparison of RKHS and RF model predictive ability and mean squared error for all phenotype clippings using the FI-2 datasets.

| Phenotype | Clipping | Predictive ability |  |  |  | Mean squared error |  |  |  |
| --- | --- | --- | --- | --- | --- | --- | --- | --- | --- |
|  |  | RKHS | RF | Diff. | Diff. (%) | RKHS | RF | Diff. | Diff. (%) |
| Green Matter | 2 | 0.857 | 0.850 | -0.007 | -0.8% | 0.018 | 0.018 | 0 | 0.0% |
|  | 3 | 0.870 | 0.869 | -0.001 | -0.1% | 0.016 | 0.017 | 0.001 | 6.3% |
|  | 4 | 0.894 | 0.912 | 0.018 | 2.0% | 0.015 | 0.013 | -0.002 | -13.3% |
|  | 5 | 0.866 | 0.917 | 0.051 | 5.9% | 0.016 | 0.012 | -0.004 | -25.0% |
|  | 6 | 0.863 | 0.903 | 0.040 | 4.6% | 0.018 | 0.012 | -0.006 | -33.3% |
|  | 7 | 0.844 | 0.923 | 0.079 | 9.4% | 0.019 | 0.010 | -0.009 | -47.4% |
|  | 8 | 0.881 | 0.913 | 0.032 | 3.6% | 0.020 | 0.015 | -0.005 | -25.0% |
|  | 9 | 0.897 | 0.944 | 0.047 | 5.2% | 0.012 | 0.008 | -0.004 | -33.3% |
|  | T | 0.867 | 0.937 | 0.070 | 8.1% | 0.018 | 0.009 | -0.009 | -50.0% |
|  | Mean | 0.871 | 0.908 | 0.037 | 4.2% | 0.017 | 0.013 | -0.004 | -24.6% |
| Regrowth | 2 | 0.868 | 0.896 | 0.028 | 3.2% | 0.015 | 0.014 | -0.001 | -6.7% |
|  | 3 | 0.940 | 0.956 | 0.016 | 1.7% | 0.011 | 0.008 | -0.003 | -27.3% |
|  | 4 | 0.859 | 0.911 | 0.052 | 6.1% | 0.017 | 0.012 | -0.005 | -29.4% |
|  | 5 | 0.826 | 0.869 | 0.043 | 5.2% | 0.016 | 0.013 | -0.003 | -18.8% |
|  | 6 | 0.833 | 0.884 | 0.051 | 6.1% | 0.024 | 0.016 | -0.008 | -33.3% |
|  | 7 | 0.862 | 0.896 | 0.034 | 3.9% | 0.015 | 0.011 | -0.004 | -26.7% |
|  | 8 | 0.860 | 0.872 | 0.012 | 1.4% | 0.014 | 0.013 | -0.001 | -7.1% |
|  | 9 | 0.813 | 0.843 | 0.030 | 3.7% | 0.017 | 0.014 | -0.003 | -17.6% |
|  | T | 0.886 | 0.932 | 0.046 | 5.2% | 0.013 | 0.008 | -0.005 | -38.5% |
|  | Mean | 0.861 | 0.895 | 0.035 | 4.1% | 0.016 | 0.012 | -0.004 | -22.8% |
| Dry Matter | 2 | 0.537 | 0.638 | 0.101 | 18.8% | 0.044 | 0.035 | -0.009 | -20.5% |
|  | 3 | 0.802 | 0.840 | 0.038 | 4.7% | 0.014 | 0.013 | -0.001 | -7.1% |
|  | 4 | 0.935 | 0.941 | 0.006 | 0.6% | 0.010 | 0.009 | -0.001 | -10.0% |
|  | 5 | 0.882 | 0.895 | 0.013 | 1.5% | 0.014 | 0.015 | 0.001 | 7.1% |
|  | 6 | 0.884 | 0.917 | 0.033 | 3.7% | 0.016 | 0.011 | -0.005 | -31.3% |
|  | 7 | 0.849 | 0.913 | 0.064 | 7.5% | 0.017 | 0.011 | -0.006 | -35.3% |
|  | 8 | 0.911 | 0.915 | 0.004 | 0.4% | 0.016 | 0.016 | 0 | 0.0% |
|  | 9 | 0.787 | 0.925 | 0.138 | 17.5% | 0.022 | 0.011 | -0.011 | -50.0% |
|  | T | 0.912 | 0.943 | 0.031 | 3.4% | 0.013 | 0.009 | -0.004 | -30.8% |
|  | Mean | 0.833 | 0.881 | 0.048 | 6.5% | 0.018 | 0.014 | -0.004 | -19.8% |
| Leaf Dry Matter | 2 | 0.835 | 0.875 | 0.040 | 4.8% | 0.019 | 0.014 | -0.005 | -26.3% |
|  | 5 | 0.899 | 0.924 | 0.025 | 2.8% | 0.012 | 0.011 | -0.001 | -8.3% |
|  | T | 0.900 | 0.952 | 0.052 | 5.8% | 0.011 | 0.007 | -0.004 | -36.4% |
|  | Mean | 0.878 | 0.917 | 0.039 | 4.5% | 0.014 | 0.011 | -0.003 | -23.7% |
| Stem Dry Matter | 2 | 0.664 | 0.818 | 0.154 | 23.2% | 0.033 | 0.021 | -0.012 | -36.4% |
|  | 5 | 0.904 | 0.921 | 0.017 | 1.9% | 0.011 | 0.012 | 0.001 | 9.1% |
|  | T | 0.694 | 0.835 | 0.141 | 20.3% | 0.025 | 0.015 | -0.010 | -40.0% |
|  | Mean | 0.754 | 0.858 | 0.104 | 15.1% | 0.023 | 0.016 | -0.007 | -22.4% |
|  | Overall Mean | 0.839 | 0.892 | 0.052 | 6.9% | 0.018 | 0.013 | -0.004 | -22.6% |

### 3.3 Major importance markers

Given that the FS strategies employed in our study relied on ML algorithms estimated through a combination of decision trees, and that the top-performing models for FI-1 and FI-2 were AB and RF, respectively, we employed an additional approach to assess marker‒trait associations using decision tree structures. We ranked the markers based on RF scores obtained from the FI-2 selected markers. We selected the top three Gini importance markers for each trait, and if the sum of importance for these top three markers did not reach at least 0.5 (out of a total of 1.0), we continued selecting markers from the ranking until we reached half of the total importance score. This process allowed us to compile a list of markers with the highest feature importance, thus preventing underrepresentation of importance across traits. From the 283 FI-selected markers across the 33 traits, we identified a subset of 69 markers with significant predictive relevance. Notably, only for SDM clipping 5, we had to select four markers instead of three (Supplementary Table S8).

Furthermore, we performed a PCA to evaluate the subfamily dispersion considering this set of 69 major importance markers. The first two PCs explained 67.5% of the data variance, an intermediate value between the complete set of SNPs (57.2%) and the phenotypic data (85.2%) (Supplementary Fig. S6). Although the values of the first PCs seem to be inverted in such a PCA when compared to the others performed, we observed a similar dispersion pattern (Supplementary Figs. S4 and S5). As we expected, the scatter plot displayed a group formation visually closer to the phenotypic PCA. Since the markers were selected through associations with the traits, there was a strong relation between the major importance data PC1 and the samples colored using the phenotypic PC1 values (Supplementary Fig. S7).

To assess the physical distribution of the FS-selected markers, we constructed a physical map for *U. ruziziensis* using the values obtained from the species’ genome. In addition to the set of 69 major importance markers, we incorporated all the FI-2 markers into the constructed map (Fig. 2). Regarding the distribution of these markers, we observed associations across all chromosomes without a clear pattern, except for the presence of extensive regions with little or no markers, primarily located in the central regions of chromosomes 1, 2, 5, 7 and 8. We speculate that these regions correspond to the centromeric regions (Fig. 2). Chromosome 1 presented the highest number of associations when considering both FI and major importance marker sets, with relatively consistent representativeness. However, it was also the chromosome with the highest number of identified SNPs (Table 2). On the other hand, chromosomes 5 and 6 presented the lowest presence of associations, while chromosome 4 experienced a significant change in representativeness, with a 7% reduction from FI to major importance (Table 2). Furthermore, especially in chromosomes 1, 4 and 7, we observed regions characterized by a high density of minor importance markers near major importance markers, which may suggest the presence of QTL regions associated with agronomic traits.

**Figure 2.**
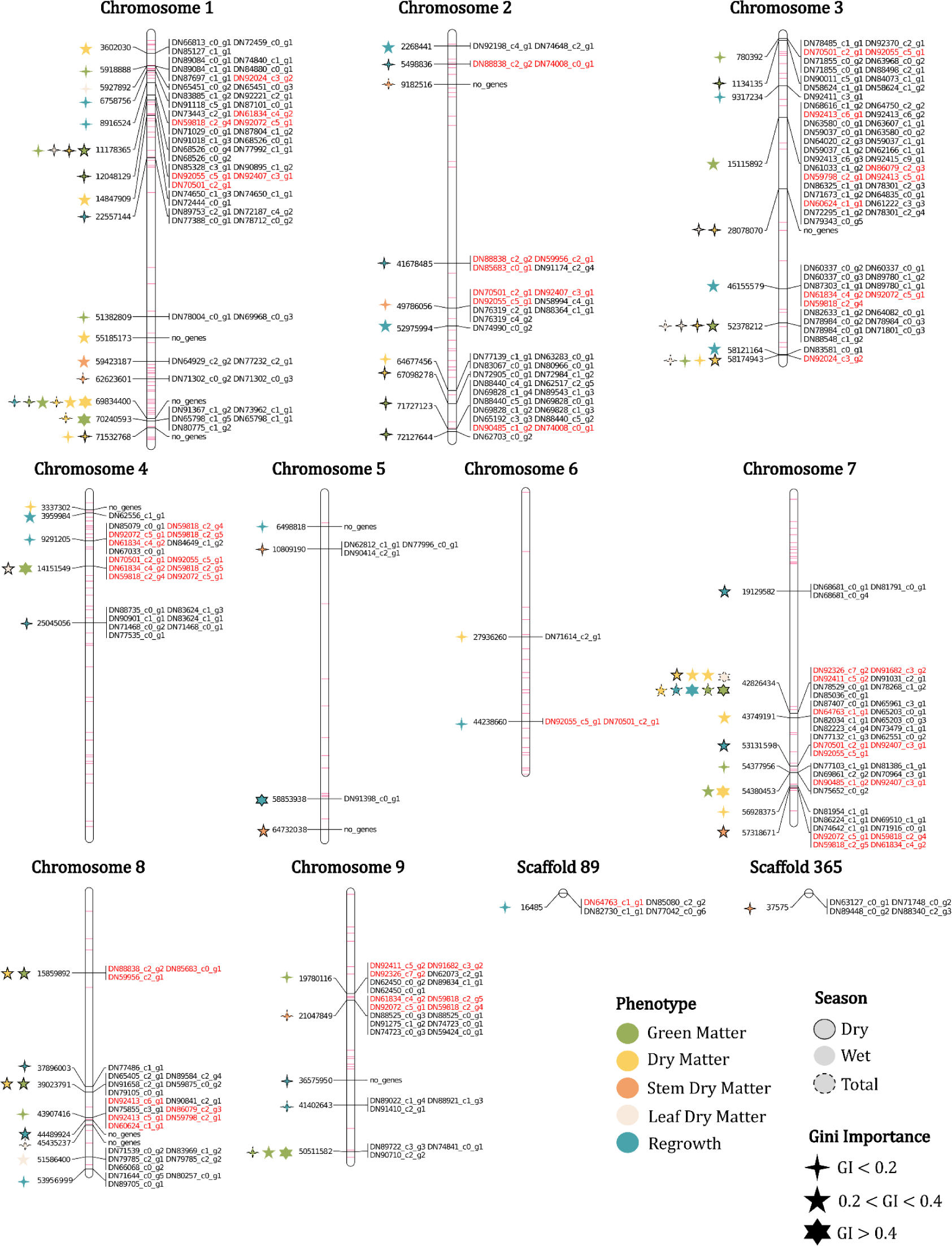
Physical map with the markers and genes associated with the phenotypes evaluated in the *U. ruziziensis* population, with Gini importance (GI) and season indicated. Duplicated genes and minor importance markers (FI-2) mapped are represented in red and purple, respectively.

**Table 2.** Number and percentage of SNP markers identified/selected in each chromosome considering the complete data (CD), feature intersection (FI-2) and top Gini importance datasets.

| Chromosome | Complete Data | Feature Intersection - 2 | Top Gini Importance |
| --- | --- | --- | --- |
| 1 | 4722 (16.8%) | 69 (23.4%) | 16 (23.2%) |
| 2 | 3565 (12.7%) | 36 (12.2%) | 10 (14.5%) |
| 3 | 3249 (11.6%) | 28 (9.5%) | 9 (13%) |
| 4 | 3552 (12.6%) | 42 (14.2%) | 5 (7.2%) |
| **5** | 1384 (4.9%) | 15 (5.1%) | 4 (5.8%) |
| **6** | 2508 (8.9%) | 23 (7.8%) | 2 (2.9%) |
| **7** | 3796 (13.5%) | 37 (12.5%) | 8 (11.6%) |
| **8** | 2127 (7.6%) | 18 (6.1%) | 8 (11.6%) |
| **9** | 2426 (8.6%) | 22 (7.5%) | 5 (7.2%) |
| **Scaffolds** | 777 (2.8%) | 5 (1,7%) | 2 (2.9%) |
| **Total** | 28106 (100%) | 295 (100%) | 69 (100%) |

The major importance set was composed of various markers associated with more than one trait. As evidenced in the physical map, the marker associated with more trait clippings is on chromosome 7, position 42,826,434. This marker was associated with four of the five phenotypes evaluated and was selected for nine clippings, three of which had a Gini importance higher than 0.4 and in six Gini importance between 0.2 and 0.4 (Fig. 2). Other markers were associated with various trait clippings, such as a marker on chromosome 1 position 69,834,400, which was associated with six trait clippings, and three other markers with four associations in chromosomes 1 and 3 (Fig. 2).

When evaluating the markers for each of the five traits without separating them by clippings, we analyzed the intersections of sets to quantify markers associated with multiple traits (Supplementary Fig. S7). Despite the variation in marker quantities between the FI-2 and major importance sets, the logical relationships among the trait sets remained consistent: GM, DM, and LDM shared the highest number of markers, while RG and SDM had a higher proportion of exclusive markers. Interestingly, SDM and RG exhibited generally lower correlations with the other traits as well.

### 3.4 Marker genes associated with phenotypes

To obtain a set of genes expressed by the species and subsequently assess their coexpression, we employed a previously published transcriptome of 11 *U. ruziziensis* genotypes. The sequencing of the libraries produced a total of ∼1.7 billion reads, with 95.5% (Supplementary Table S9) being retained and used for de novo assembly. The resulting transcriptome encompassed 575,524 transcripts, of which 223,593 were categorized as unigenes, featuring a transcript N50 length of 1,227 bp. Following filtration based on expression levels, the dataset was reduced to 288,487 transcripts, representing 49,445 unigenes. The evaluation of assembly completeness was performed by comparing the 49,445 unigenes against the Viridiplantae database. From the 425 total BUSCO groups searched, we found 297 complete sequences (69.8%), 48.2% as a single copy and 21.6% as duplicated copies, in addition to 74 (17.4%) and 54 (12.8%) fragmented and missing sequences, respectively.

In the process of functional annotation, we aligned the transcripts to the UniProt database and obtained 197,045 associated GO terms. Among these, 6,156 were unique GO terms. This collection of genes and GO terms was then employed to perform a biological process GO term enrichment analysis of the genes linked to the major importance markers.

After aligning transcripts with the reference genome of *U. ruziziensis* and considering a window of 5,000 bp up/downstream of the marker positions, we mapped a total of 217 genes (264 considering genes with multiple copies) in close physical proximity to 58 markers (Fig. 2 and Supplementary Table S8). We did not detect genes linked to all markers, such as on chromosome 1, where no genes were found within a marker region associated with six traits clippings, or on chromosome 5, where out of the four major importance markers, two lacked associated genes (Fig. 2).

As previously stated, we identified genes with multiple copies that are linked to more than one major importance marker region. There were 22 genes meeting this criterion, and they are highlighted in red in Fig. 2. To facilitate a more comprehensive investigation of these genes, we represented their distribution in a circular map that illustrates their genomic positions. Additionally, we combined information about copy number variation, trait/season associations, and Gini importance (Fig. 3). Among these genes, we identified five genes with six copies. Notably, three of these genes are found together, and collectively, they are associated with seven different trait categories. Furthermore, we identified genes with 4, 3 and 2 copies, all linked to all the evaluated traits, albeit with varying levels of importance, demonstrating no clear pattern.

**Figure 3.**
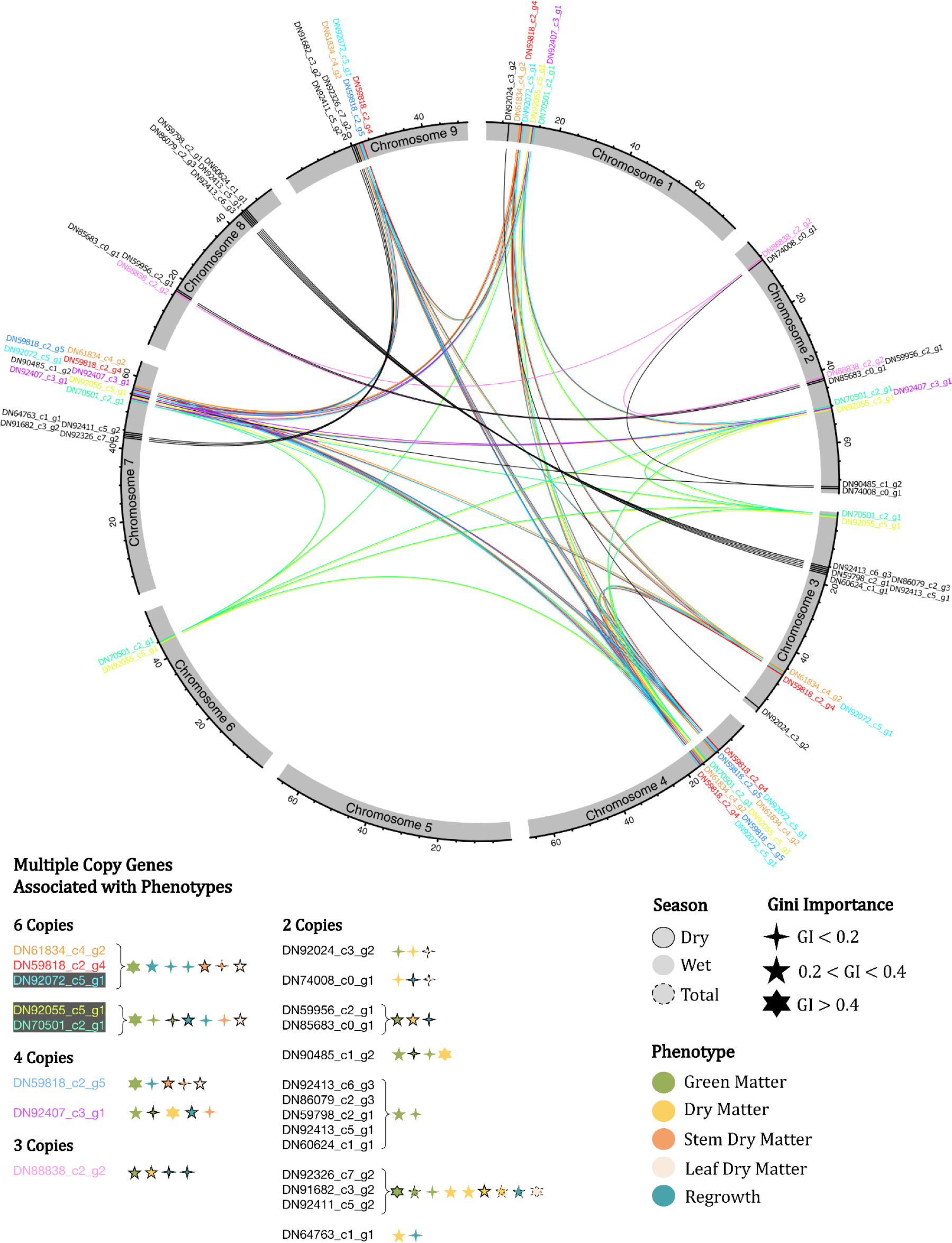
Circular map of the *U. ruziziensis* genome, indicating multiple copy genes identified as associated with the phenotypes evaluated, indicating Gini importance and season. The same genes are indicated with the same color, except genes with two copies.

Regarding the functional annotation of the genes associated with the phenotypes, we identified proteins/enzymes and GO terms for 100 of the 217 genes (Supplementary Table S8). In the region associated with more traits, on chromosome 7 (position 42,826,434), seven annotated genes were mapped, some of which were cinnamoyl-CoA reductase 1, and DEAD-box ATP-dependent RNA helicase 25. Furthermore, on chromosome 1 (position 11,178,365), which is associated with four traits, there are genes annotated to the multidomain protein RHM2/MUM4 which is involved in UDP-D-glucose to UDP-L-rhamnose conversion (Supplementary Table S8). Considering the genes with multiple copies, only three had functional annotation, which translates into AIM25-altered inheritance rate of mitochondria protein 25, cinnamoyl-CoA reductase 1 and E3 ubiquitin-protein ligase SINAT5.

Beyond specific protein annotation, to obtain a general functional profile of the proteins identified, we performed an enrichment analysis of the biological process GO terms and obtained a profile with 18 significant terms (p value < 0.01). The enrichment analysis identified terms associated with various phenotype clippings, such as “lignin biosynthetic process”, “auxin efflux” and “flavonol biosynthetic process” (Supplementary Table S10).

### 3.5 Coexpression network

To provide deeper insights into the functional patterns of genes associated with the agronomic traits evaluated, we modeled a GCN using the gene quantifications from the *U. ruziziensis* accessions. From a total of 49,445 genes, we identified significant interactions between 14,141 genes, represented as nodes in the network structure, connected by 17,812 edges (Supplementary Fig. S8). Within this GCN, we found 54 genes from the 217 genes associated with the major importance markers. As we restricted the GCN created to the top 30 gene associations, we expanded the collection of 54 selected genes to more than 109 by considering correlations with a minimum Pearson coefficient of 0.9 and a Bonferroni corrected p value of 0.01. This group of 153 genes was considered directly associated with the traits evaluated.

The potential of a GCN to elucidate metabolic pathways lies in its ability to identify genes that, despite not being selected by the prediction methodology, exhibit coexpression with them. To this end, we extended the set of 153 genes previously selected to the GCN first (308 genes) and second gene neighbors (2233 genes), creating a comprehensive agronomic trait network comprising a total of 2704 genes (nodes) and 3453 edges (Fig 4).

**Figure 4.**
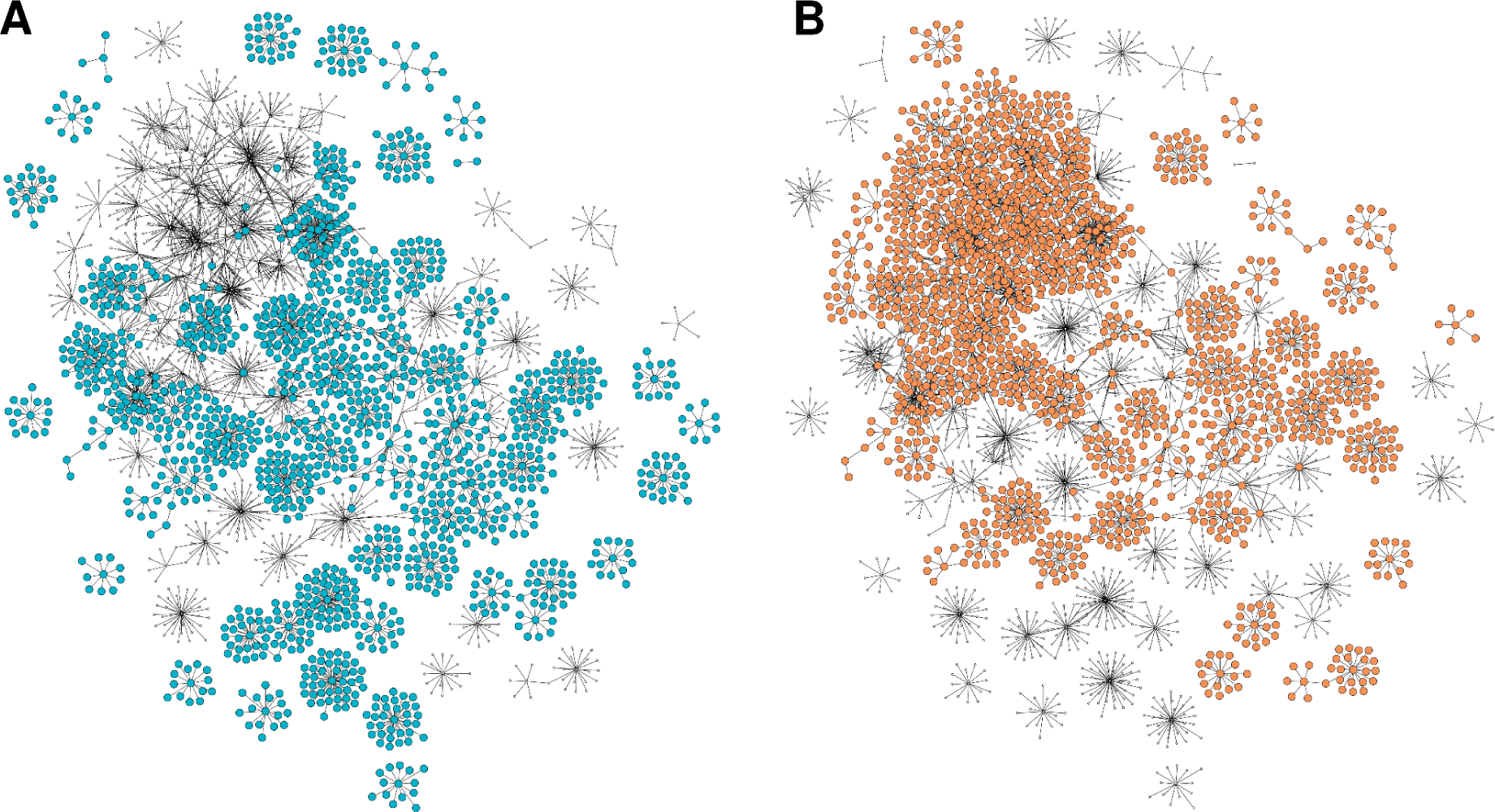
Selected and correlated gene coexpression network with first and second neighbors. A) Major importance genes in blue, highly correlated genes in yellow, first neighbors in green and second neighbors in gray. B) Genes associated with wet season trait clippings. C) Genes associated with dry season trait clippings.

The functional profile of the general agronomic traits network was determined through an enrichment analysis of biological process GO terms, which revealed 11 significant terms (p < 0.01) for the gene set excluding the second neighbors and 16 terms when considering all genes in the network (Supplementary Table S11). When examining the restricted set, which excluded the second neighbors, we found enriched terms related to hormones, such as auxin efflux and abscisic acid transport, as well as biosynthetic processes involving molecules such as flavonoids. In the broader set that included second neighbors, we identified terms associated with DNA metabolism, including mismatch repair, DNA replication, and DNA duplex unwinding. Additionally, other enriched terms were linked to responses to stress, such as response to chitin and regulation of circadian rhythm.

To further explore the differences in functional gene patterns associated with the different seasons, we separated the general agronomic trait network into two seasonal parts. The genes associated with the traits in clippings 2,3,7,8 and 9 were selected for the wet season-associated network, and the genes associated with the traits in clippings 4, 5 and 6 were selected for the dry season-associated network. The wet and dry-season networks encompassed 33 and 22 genes associated with the major importance markers, 58 and 54 highly correlated genes, 102 and 231 first neighbors, 1322 and 1359 second neighbors, and a total of 1515 and 1666 genes (nodes) with 1801 and 2205 edges, respectively (Fig 4-A, B). Comparing the seasonal functional profiles, we found shared terms such as flavonol biosynthetic process, auxin efflux and mitotic recombination-dependent replication fork processing. Additionally, we discovered season-specific terms such as abscisic acid transport, isoleucine biosynthetic process, response to nematode and chaperone-mediated protein folding for the wet season. In contrast, the dry season featured enriched terms such as pyridoxal phosphate biosynthetic process, response to water deprivation and response to chitin, all of which are related to stress response (Supplementary Table S12 and S13).

Another remarkable aspect of using GCNs to investigate the regulation of metabolic pathways lies in their ability to define hub genes, which possess a high number of connections in the network, as determined by the degree metric. The hub genes play an important role in regulating the functionality of numerous other genes, thereby potentially influencing the expression of the phenotypes that we are studying. In our modeled agronomic traits network, we found 235 hub genes (degree > 4), of which 14 had a degree > 40. Considering the seasonal networks, there were 107 and 158 hubs in the wet and dry season networks, respectively. Among the highest degree hub genes (>40), we found some present in both season networks, such as the genes that encode the 60S ribosomal protein L9 and the 14-3-3 protein zeta (Supplementary Table S14). While specific to the wet season, we found hub genes of the proteins ELF4-LIKE 4, SUV2 and lipid-transfer DIR1 (Supplementary Table S15) and to the dry season, 60S ribosomal L9, fatty acid-binding and 3-hydroxyacyl dehydratase FabZ (Supplementary Table S16).

## 4 Discussion

### 4.1 Genome-wide family prediction

Recent advances in omics approaches and computational methods for polyploid species have enabled the emergence of studies in important *Urochloa* breeding areas. These include genome assembly (Pessoa-Filho et al., 2019; Worthington et al., 2021), contaminant identification (Martins et al., 2021), transcriptomics (Vigna et al., 2016a; Salgado et al., 2017; Worthington et al., 2021; Jones et al., 2021; Hanley et al., 2021), linkage and QTL mapping (Vigna et al., 2016b; Thaikua et al., 2016; Ferreira et al., 2016; Worthington et al., 2016; Worthington et al., 2019; Worthington et al., 2021), GWAS (Matias et al., 2019b), and GS/GP (Matias et al., 2019a; Aono et al., 2022). Even with the recent progress, there are no studies employing integrative methodologies in the genus. Therefore, by leveraging the limited genetic resources available and employing biocomputational techniques, we have pioneered the first multiomic approach in the *Urochloa* genus, thereby expanding the molecular knowledge available for breeding.

In our initial approach to evaluating GWFP, we employed the complete marker dataset (CD) with conventional parametric and semiparametric GP models (BRR and RKHS). Our findings consistently yielded either higher or, at the very least, equivalent PA when compared to the achievements reported in other GWFP studies. While we achieved a high mean PA of ∼0.8 for the DM clippings, in the case of alfalfa, the authors observed values below 0.7 for the same trait in both 10-fold and leave-one-group-out cross-validation scenarios (Murad Leite Andrade et al., 2022). Additionally, for ryegrass, an even lower PA value of 0.34 was observed in a leave-one-family-out cross-validation scenario (Guo et al., 2018). If we consider the GWFP results for other phenotypes, such as rust resistance and heading date in alfalfa (Murad Leite Andrade et al., 2022), as well as lignin content, stiffness and diameter in loblolly pine (Rios et al., 2021), they consistently exhibited smaller PA values compared to our results, which presented a mean PA of ∼0.762 across the 33 traits. We attribute this PA in our predictions primarily to the relatively small population size and limited genetic diversity among our samples. This combination has been previously reported to enhance predictive accuracy, as demonstrated in wheat (Edwards et al., 2019). Moreover, increasing the population with genetically distant samples tends to increase the complexity of the prediction task, subsequently reducing its accuracy, as highlighted in previous studies (Lorenz & Smith, 2015; Berro et al., 2019).

The performance we achieved seems even more promising when compared to GP conducted at the individual level in tropical forages. Studies with *U.ruziziensis* interspecific hybrids (Matias et al., 2019a), *U. decumbens* (Aono et al., 2022), *M. maximum* (de C. Lara et al., 2019; Aono et al., 2022), and *P. virgatum* (Lipka et al., 2014) have yielded PAs ranging from values close to zero to maximum values of 0.7 when evaluating several agronomic and morphological traits and employing different cross-validation schemes. In addition to the well-known advantages of genomic selection, such as its potential to shorten the breeding process (Simeão-Resende et al., 2014) and reduce phenotyping costs (Crossa et al., 2017), the use of GWFP offers other advantages, particularly in forage breeding programs, which typically rely on family-or plot-level phenotyping for conventional breeding (Rios et al., 2021). Furthermore, when GWFP is conducted using ML models combined with FS/FI strategies, it has the potential to significantly lower genotyping costs.

Moreover, for the application of GWFP in breeding programs or for future research, we recommend prioritizing experimental designs featuring a greater number of families rather than increasing the number of individuals within each family. Studies with tetraploid full-sibling families have concluded that using six individuals is sufficient to effectively capture family variation, both in terms of genotyping and phenotyping (de Bem Oliveira et al. 2020; Rios et al., 2021). Furthermore, it is worth noting that to a certain extent, enlarging the training population holds the potential to enhance the performance of GWFP, as indicated by previous research (Fé et al. 2015).

The application of ML algorithms in GP has been extensively explored across various species and phenotypes (Grinberg et al., 2016; Lello et al., 2018; Zhao et al., 2020; Liang et al., 2020; Chung et al., 2021; Islam et al., 2021; Sandhu et al., 2021). Although there is no concrete empirical evidence supporting the superiority of ML over linear methods (Zingaretti et al., 2020; Varshney, 2021), ML techniques have consistently demonstrated superior or at least equivalent performance compared to conventional models in diverse scenarios (Bellot et al., 2018; Abdollahi-Arpanahi et al., 2020; Liang et al., 2021; Wang et al. 2022). ML methods have the potential to outperform conventional GP models, especially when handling intricate phenotypes influenced by significant dominance and epistatic effects (Wang et al., 2018; Tong & Nikoloski, 2021). Moreover, there are no studies investigating the applicability of ML methods in GWFP. Thus, we evaluated four classical ML algorithms (SVM, RF, AB and MLP). Surprisingly, none of these algorithms was able to outperform the RKHS model. Although SVM demonstrated competitive performances for PA, it was not as good for MSE. RF and AB exhibited intermediate performances, whereas MLP markedly underperformed (Supplementary Tables S2-4). The poor performance of MLP may be attributed to the limited sample size of our dataset and the lack of hyperparameter tuning. Neural network methods are well known for their need for substantial datasets and meticulous hyperparameter tuning to achieve high prediction accuracy (Bellot et al., 2018; Montesinos-López et al., 2021).

### 4.2 Major Importance Markers

Our study goes beyond the applicability analysis of GWFP in *U. ruziziensis*. We also aimed to investigate the metabolic regulation of agronomic traits. As an initial step to achieve this objective, we aimed to establish potential marker-phenotype associations. To this end, the strong performance improvement observed using the FS/FI approach indicates that the selected sets of markers are likely to be near QTLs, and can therefore be used to define genomic regions involved in phenotypic variation (Steinfath et al., 2010; Heer et al., 2018; Zhou et al., 2019; Aono et al., 2020; Pimenta et al., 2021; Aono et al., 2022; Pimenta et al., 2022). In contrast to other approaches aimed at identifying genotype-phenotype associations, FS techniques do not rely on specific biparental populations (RILs, NILs, F2, etc.), which are necessary for QTL mapping (Mohan et al., 1997; Dhingani et al., 2015). Moreover, FS techniques have the ability to uncover nonlinear and complex associations, addressing a limitation of linear models used in GWAS (Korte & Farlo, 2013).

ML models based on decision trees offer good prediction interpretability since it is possible to assess feature importance. In the context of GP, these models can rank markers based on their association strength with the modeled phenotype (Azodi et al., 2019b; Bayer et al., 2021; Medina et al., 2021). Therefore, given that the best model for each FI dataset type was equivalent, we computed the RF Gini importance for the more restricted FI-2 datasets and selected the most significant features to create an even smaller and more reliable set of putatively agronomic trait-associated markers. By using half-sibling families’ bulks as a representation of the genetic variability available for breeding, genotyping similar agronomic traits in various clippings and selecting only the most influential markers in the predictions, we were able to minimize the limitations of the method due to small sample size and obtain a reliable set of markers In this major importance set, we identified markers associated with multiple phenotypes. Notably, the number of shared markers was more pronounced for GM, DM and LDM, which is in accordance with the observed correlations among phenotype clippings, where SDM and RG exhibited lower correlations with the other traits (Supplementary Fig S3 and S6). The high overall correlation, with a mean of 0.72, among traits and the overlap of the identified markers were as expected. This is because the assessed biomass characteristics are highly similar and likely influenced by the same metabolic processes. The DM phenotype was determined by drying the GM material, while SDM/LDM phenotyping involved separating the DM into stems and leaves. Furthermore, biomass production is dependent on the plant’s growth capacity (RG). Consequently, the narrow-sense heritabilities of these traits within the families were also very similar. As discussed in other studies, modeling performance is strongly influenced by heritability (Wang et al., 2018; Xu et al., 2018; Murad Leite Andrade et al., 2022). Therefore, our prediction performances did not vary significantly and were correlated with the heritabilities (Supplementary Tables S1, S2 and S3).

In the absence of genome annotation, we employed RNA-Seq data in a multiomic approach to map genes physically associated with the major importance markers. Considering the similarity of the agronomic traits employed and the potential involvement of the same biological processes in their regulation, we then conducted a functional analysis that considered the annotation of all genes collectively. This allowed for an overview of the most influential processes governing biomass production and growth.

The enrichment of GO terms related to the annotated genes provides evidence of the methodological capacity to identify QTL regions influencing the evaluated agronomic traits (Supplementary Table S10). Associated with various phenotype clippings, terms related to the lignin biosynthetic process stood out. Previous research has established its significant impact on plant development (Yoon, Choi & An, 2015; Bahri et al., 2020). Mutants of lignin biosynthesis genes have shown phenotypes of dwarfism/reduced plant growth (Schilmiller et al., 2009; Wagner et al., 2009; Li et al., 2009; Song & Wang, 2011), altered morphology (Elkind et al., 1990; Piquemal et al., 1998; Jones et al., 2001; Franke et al., 2002), and tissue browning (Bout & Vermerris, 2003; Xu et al., 2011; Saballos et al. 2012). Furthermore, terms associated with auxin efflux were identified, which are known for their importance in growth regulation. Auxin hormone effects depend on concentration and are primarily produced in meristematic and specific regions (Blakeslee, Peer & Murphy, 2005; Zhao, 2018). The transport and distribution of auxin within plant tissues constitutes an essential aspect of its function in plant organogenesis and morphogenesis (Woodward & Bartel, 2005). This transport is facilitated by influx and efflux carrier proteins, providing essential directional and positional cues for various developmental processes, including vascular differentiation, apical dominance, organ development, and tropic growth (Benkova et al., 2003; Blancaflor & Masson, 2003; Friml et al., 2003; Blilou et al., 2005; Grieneisen et al., 2007).

Furthermore, the flavonol biosynthetic process, which is another enriched term identified in our results, is known to regulate plant growth and development by controlling auxin transport. Its effects are primarily observed in root elongation, quantity and gravitropic response (Jacobs & Rubery, 1988; Brown et al., 2001; Santelia et al., 2008; Grunewald et al., 2012). Flavonols can influence auxin transportation by different mechanisms, such as modulating the transcription of genes encoding auxin transport proteins (Peer et al., 2004), acting as kinase inhibitors that regulate the phosphorylation status of auxin transport proteins (Agullo et al., 1997; Peer & Murphy, 2007), or altering the cellular redox state (Fernández-Marcos et al., 2013). In addition, flavonols have antioxidant functions, acting in response to stress such as UV radiation, wounding, drought, metal toxicity, and nutrient deprivation. These conditions lead to the accumulation of reactive oxygen species (ROS), which can damage cellular components and consequently impact plant development (Winkel-Shirley, 2001; Baskar, Venkatesh & Ramalingam, 2018; Agati et al., 2020). The list of terms associated with plant growth, development and stress response continues with the folic acid biosynthetic process (Stakhove et al., 2000; Gorelova et al., 2017), galactolipid metabolic process (Kobayashi et al., 2007; Jouhet et al., 2007; Botté et al., 2011), and cellular response to cold.

The enriched terms provided an overview of the biological function of the identified genes. However, for the genes with multiple copies, limited information was generated, as only three out of the 22 genes had functional annotation. Nevertheless, these three genes appear to have a significant impact on the evaluated agronomic traits. One of these genes, DN91682_c3_g2 (cinnamoyl-CoA reductase 1), which was identified in two copies, is involved in the lignin biosynthetic process (Lauvergeat et al., 2001), circadian rhythm, and response to cold (Carpenter et al., 1994). The second gene, DN64763_c1_g1 (E3 ubiquitin-protein ligase SINAT5), also found in two copies, is known to play key roles in multiple plant developmental stages and several abiotic stress responses (Shu & Yang, 2017). Furthermore, although it has been reported in yeast, the gene DN92072_c5_g1 (AIM25-altered inheritance rate of mitochondria protein 25), which was found in six copies linked to major importance markers, acts in the cellular response to heat and oxidative stress (Aguilar-Lopez et al., 2016) (Fig. 3) (Supplementary Table S8).

As a result of diverse mechanisms, such as whole-genome duplication, tandem duplication, and transposon-mediated duplication, plant genomes have an abundance of duplicated genes (Panchy, Lehti-Shiu & Shiu, 2016). These duplicate copies can persist for several reasons: insufficient time for the accumulation of deleterious mutations or selection pressure to preserve redundant functions (Panchy, Lehti-Shiu & Shiu, 2016). This pressure can arise from four mechanisms: gene dosage increase (Ohno, 1970), subfunctionalization (Force et al., 2013), gene balance (Freeling & Thomas, 2006), and paralog interference (Baker et al., 2013). Beyond identifying multiple copies of genes associated with agronomic traits, further investigation into the mechanisms influencing their retention and how these copies interact and impact the trait may provide valuable insights for improving breeding methods to achieve higher genetic gains.

### 4.3 Gene Coexpression Network

We conducted additional multiomic investigations to gain a deeper understanding of the metabolic pathways and regulatory mechanisms that govern the evaluated agronomic traits. We modeled a GCN and isolated the previously identified genes and their coexpressed neighbors (Fig. 4-A). This integration has been successfully employed in different species and has produced noteworthy results (Calabrese et al., 2017; Schaefer et al., 2018; Yan et al., 2020; Francisco et al., 2021). The ability of such networks to simulate complex biological systems and uncover novel biological associations has transformed molecular biology research (D’haeseleer et al., 2000; Liu et al., 2020), enabling the exploration of regulatory relationships, inference of metabolic pathways, and transfer of annotations (Rao & Dixon, 2019). Following the “guilt-by association” principle, GCNs typically reveal interactions among genes with correlated biological functions (Oliver, 2000, Wolfe et al., 2005, Childs et al., 2011).

In this context, we could expand our set of identified genes through coexpression analysis, providing broader insight into the metabolic pathways influencing the observed phenotypes. Moreover, the annotated genes within these modules can serve as a basis to infer the biological functions of the unannotated genes. Our network modeling has extended our understanding of genes associated with the previously discussed enriched terms. It has also enabled the identification of new genes involved in biological processes related to DNA integrity, stability and metabolism. These genes act in mismatch repair, telomere capping, and duplex unwinding, all of which are known to impact the normal growth and development of plants to varying degrees (Karthika et al., 2020, Kim & Kim, 2018, Tuteja, 2003). Additionally, our network also expanded the genes involved in regulating the abscisic acid (ABA) transport. Modulating hormone levels within tissues and cells is critical for maintaining a balance between defense mechanisms and growth processes, especially in suboptimal environments. This regulation also plays an important role in controlling stomatal closure (Seo & Koshiba, 2011; Chen et al., 2020). Furthermore, the network has elucidated genes involved in regulating the circadian rhythm. Such a process not only allows plants to adapt to daily environmental changes but also enables them to anticipate and prepare for these challenges in advance (Kim et al., 2017; Millar, 2016; Creux & Harmer, 2019). Notably, the gene ELF4-LIKE 4, a key player in the circadian rhythm (Doyle et al., 2002), stands out as one of the hub genes with the highest degree value in the network (Supplementary Table S12). Finally, we also identified genes related to response to chitin, an important component of the plant immune system activated in the presence of pathogens such as fungi, arthropods, and nematode egg shells (Kombrink et al., 2011; Sánchex-Vallet et al., 2015).

Furthermore, by separating the general agronomic trait network into two seasonal parts, we were able to investigate the potential impact of metabolic processes on plant development and production during both wet and dry periods. In our findings, we identified enriched terms related to auxin efflux and flavonol biosynthetic processes in both networks. These results have already been discussed in the context of auxin transport regulation, indicating the importance of the hormone regardless of the season. During the wet season, in addition to the previously mentioned abscisic acid transport, we observed terms associated with plant development, such as the isoleucine biosynthetic process (Yu et al., 2013) and the response to nematodes, which are pathogens capable of modifying plant physiology, development, metabolism, and immunity (Eves-van den Akker, 2021). In contrast, within the dry network, we found enriched terms related to responses to water deprivation and protein transport. This provides evidence of the metabolic mechanisms required to deal with abiotic stress.

Network degree analysis provided a means to identify hub genes, which are the most highly connected genes in the network. These hubs typically encompass genes with broad regulatory functions or associations with essential roles in biological processes (Carlson et al., 2006; Reverter et al., 2008; Amrine et al., 2015). In our analysis, in addition to the previously mentioned ELF4-LIKE 4 protein, we identified several ribosomal protein genes as hubs, such as 40S S6 and S15a-2, 60S L9 and L14-2, 54S L12, and Ubiquitin-40S S27a-1. The heterogeneity of ribosome composition is well-known and forms the foundation of the specialized ribosome theory, which states that different groups of ribosomes are tailored to translate specific sets of mRNAs (Gilbert, 2011; Xue & Barna, 2012, Genuth & Barna, 2018). Although the major discussion in this field is concentrated in elucidating how changes in ribosome composition might facilitate the translation of specific groups of mRNAs (Norris, Hopes & Aspden, 2021), our results indicate another intriguing aspect of this theory. Although the precise connection between the observed hub ribosomal proteins and the translation of the genes linked to the hubs has yet to be established, we hypothesize that their coexpression may result from a coregulatory mechanism that ensures the availability of specific tailored ribosomes in sufficient quantities for translating the mRNAs of these linked genes. Regarding the relationship between ribosomal proteins and the characteristics evaluated in this research, it has been reported that *A. thaliana* mutants in these proteins are often smaller and have simplified/aberrant vasculature and polarity defects (Van Lijsebettens et al., 1994; Ito et al., 2000; Fujikura et al., 2009; Horiguchi et al., 2011), which can directly impact attributes related to regrowth and biomass production.

In addition, among the highest degree hubs in the network, we found genes associated with lipid metabolism. These specific genes encode important proteins, including the lipid-transfer protein DIR1, fatty acid-binding protein, 3-hydroxyacyl-ACP dehydratase, and 3-Ketoacyl-CoA Synthase 4. These proteins play roles in fatty acid biosynthesis (Supplementary Table S14). Fatty acids, which are common components of complex lipids, are reported to have important roles in plant biology, including cell structure and response to different stresses such as temperature changes (Routaboul et al., 2000; Iba, 2002; Hou et al., 2006), salinity, drought (Mikami & Murata, 2003; Gigon et al., 2004; Zhang et al., 2005), exposure to heavy metals (Verdoni et al., 2001; Chaffai et al., 2007; Maksymiec, 2007), and pathogens (Kachroo et al., 2003; Nandi et al., 2005). Fatty acids, as integral components of cellular membranes, suberin, and cutting waxes (Beisson et al. 2007), contribute to stress resistance by modulating membrane fluidity, releasing α-linolenic acid (Grechkin 1998), serving as precursors for bioactive molecules (Hou et al., 2016), and acting as modulators of plant defense gene expression (Kachroo et al. 2001).

Another gene with broad activity identified as a hub is 2-oxoglutarate/Fe(II)-dependent dioxygenase (2-ODD) (Supplementary Table S14). This highly versatile enzyme facilitates numerous oxidative reactions, playing a crucial role in biosynthetic pathways for normal organismal function and the production of high value specialized metabolites (Farrow & Facchini, 2014). Its roles extend across various pathways, including DNA repair, histone demethylation, posttranslational modifications, auxin and salicylic acid catabolism, and biosynthesis of gibberellin, ethylene, flavonoid and glucosinolate. 2-ODD is reported to have a significant impact on plant growth and development (Farrow & Facchini, 2014).

Another important aspect of the methodology employed lies in its ability to identify regions associated with known genes linked to specific traits. Equally important is its capacity to elucidate unannotated genes that should be investigated. In our results, more than half of the identified genes linked to the major importance markers lacked functional annotation. Remarkably, some of these unannotated genes seemed to be highly important, as they were observed to have multiple copies and associations with various traits (Fig. 3). When we expanded our analysis to the GCN, even more unannotated genes emerged, including important hub genes evidenced by their high degree values (Supplementary Table S14). These genes/regions are important targets to expand the knowledge on the metabolic regulation of agronomic traits and represent valuable information that can be applied in species breeding.

Our work is innovative in different aspects and represents a significant advance in the field of molecular breeding techniques applicable to tropical forages. This study marks the first exploration of the applicability of GWFP in a *Urochloa* species, being the first time that FS and ML algorithms have been employed in GWFP. These techniques not only enhance prediction metrics but also drastically reduce the number of makers required for accurate prediction. Furthermore, employing a multiomic approach, we integrated the selected markers with transcriptome data to construct a coexpression network capable of providing insights into the regulation of plant growth and biomass production in the species. The results demonstrate the great potential of molecular breeding in reducing breeding costs, expediting the release of new cultivars, and facilitating metabolic investigations, even in orphan species with high genomic complexity, such as tropical forages.

## 6 Author Contributions

FBM, AHA, RMS and APS conceived the study. RMS conducted the field experiments. RMS, ACLM, RCUF, MMV and MPF performed the laboratory experiments. FBM and AHA analyzed the data. MRM assisted statistical data analyses. FBM, AHA, ACLM and RCUF wrote the manuscript. All the authors have read and approved the manuscript.

## Acknowledgments

We would like to acknowledge the Fundação de Amparo à Pesquisa de do Estado de São Paulo (FAPESP), the Conselho Nacional de Desenvolvimento Científico e Tecnológico (CNPq), and the Coordenação de Aperfeiçoamento de Pessoal de Nível Superior (CAPES). We also acknowledge the Brazilian Agricultural Research Corporation (Embrapa Gado de Corte) for providing the populations used in this study. This manuscript was previously posted to bioRxiv https://www.biorxiv.org/.

## 8 Data Availability statement

The raw sequence data for *U. ruziziensis* have been submitted to the NCBI Sequence Read Archive (SRA) under accession number PRJNA973612.

## Supplementary Tables

**Supplementary Table S1.** Estimates of narrow-sense heritability (h²) for all phenotypes, stratified by clippings and assessed at the individual level.

**Supplementary Table S2.** Comparison of model performances assessed by predictive ability and mean squared error, separated according to phenotype and clipping. For each specific combination of trait, clipping, and dataset configuration, a distinct Tukey’s test was conducted to compare model performance. The compared datasets encompassed the following: (i) the complete dataset, (ii) markers selected by at least two feature selection techniques (FI-1), and (iii) markers selected by all three employed feature selection techniques (FI-2). The tested models were the semiparametric reproducing kernel Hilbert space (RKHS), Bayesian ridge regression (BRR), support vector machine (SVM), random forest (RF), adaptive boosting (AB), and multilayer perceptron (MLP) neural network.

**Supplementary Table S3.** Comparison of model performances assessed by predictive ability and mean squared error, separated according to phenotype and clipping. For each specific combination of trait, clipping, and dataset configuration, a distinct Tukey’s test was conducted to compare clippings. The compared datasets encompassed the following: (i) the complete dataset, (ii) markers selected by at least two feature selection techniques (FI-1), and (iii) markers selected by all three employed feature selection techniques (FI-2). The tested models were the semiparametric reproducing kernel Hilbert space (RKHS), Bayesian ridge regression (BRR), support vector machine (SVM), random forest (RF), adaptive boosting (AB), and multilayer perceptron (MLP) neural network.

**Supplementary Table S4.** Number of scenarios identified as optimal by Tukey’s test. The total number of scenarios corresponds to the number of phenotype clippings (33). The compared datasets encompassed the following: (i) the complete dataset, (ii) markers selected by at least two feature selection techniques (FI-1), and (iii) markers selected by all three employed feature selection techniques.

**Supplementary Table S5.** Numbers of markers selected by the three feature selection (FS) methods. This evaluation was conducted for all phenotypes and clippings, along with their intersections (FI), considering markers selected by at least two feature selection techniques (FI-1) and markers selected by all three employed feature selection techniques.

**Supplementary Table S6.** Number of scenarios in which the evaluated datasets were determined as the optimal choice based on Tukey’s test. A total of 198 scenarios were evaluated, considering 6 models and 33 phenotype clippings.

**Supplementary Table S7.** Optimal scenarios identified by Tukey’s test for model, phenotype, and dataset comparison. Models highlighted in red represent the most favorable choices for the respective dataset.

**Supplementary Table S8.** Functional annotation of the transcripts physically linked to the major importance markers defined.

**Supplementary Table S9.** Count of sequencing reads pre- and postquality filtering procedures.

**Supplementary Table S10.** Biological process Gene Ontology (GO) enrichment analysis of genes physically linked to the major importance trait-associated markers.

**Supplementary Table S11.** Gene Ontology (GO) enrichment analyses of the modeled gene coexpression network considering (i) genes associated with traits and their first coexpressed neighbors and (ii) genes from (i) and their coexpressed neighbors (second neighbors from the genes associated with traits).

**Supplementary Table S12.** Gene Ontology (GO) enrichment analyses of the modeled gene coexpression network considering (i) genes associated with traits measured during the wet season and their first coexpressed neighbors and (ii) genes from (i) and their coexpressed neighbors (second neighbors from the genes associated with traits).

**Supplementary Table S13.** Gene Ontology (GO) enrichment analyses of the modeled gene coexpression network considering (i) genes associated with traits measured during the dry season and their first coexpressed neighbors and (ii) genes from (i) and their coexpressed neighbors (second neighbors from the genes associated with traits).

**Supplementary Table S14.** Degree assessment of nodes within the modeled gene coexpression network encompassing genes linked to traits along with their first and second-degree neighbors.

**Supplementary Table S15.** Degree assessment of nodes within the modeled gene coexpression network encompassing genes linked to traits measured during the wet season along with their first and second-degree neighbors.

**Supplementary Table S16**. Degree assessment of nodes within the modeled gene coexpression network encompassing genes linked to traits measured during the dry season along with their first and second-degree neighbors.

## Supplementary Figures

**Supplementary Figure S1.** Climatological water balances using the available water capacity (AWC = 100 mm) on a 10-day scale from November 2012 to January 2014 at the Brazilian Agricultural Research Corporation (Embrapa) Beef Cattle (EBC) station in Campo Grande, Mato Grosso do Sul State, Brazil (20°27’S, 54°37’W, 530 m).

**Supplementary Figure S2.** Box plots of the scaled phenotype clippings. DM, LDM, SDM, GM and RG represent dry matter, leaf dry matter, stem dry matter, green matter and regrowth, respectively.

**Supplementary Figure S3.** Heatmap showing the correlation between all trait clippings. Red lines separate different phenotypes. DM, LDM, SDM, GM and RG represent dry matter, leaf dry matter, stem dry matter, green matter and regrowth, respectively.

**Supplementary Figure S4.** Principal component analysis scatter plot of the family phenotypes. The axes represent the first and second principal components, which explain 75.7% and 9.5% of the variance, respectively.

**Supplementary Figure S5.** Principal component analysis scatter plot of family genotyping, with a total of 28,106 markers. The axes represent the first and second principal components, which explain 38.9% and 18.3% of the variance, respectively.

**Supplementary Figure S6.** Principal component analysis scatter plot of family genotyping, considering only the major importance markers (69 markers). The axes represent the first and second principal components, which explain 58% and 9.5% of the variance, respectively.

**Supplementary Figure S7.** Venn diagrams showing the logical relationship among the sets of markers identified for each phenotype. (A) A total of 283 markers from the FI-2 dataset and (B) 69 markers selected by the Gini importance condition. DM, LDM, SDM, GM and RG represent dry matter, leaf dry matter, stem dry matter, green matter and regrowth, respectively.

**Supplementary Figure S8.** *Urochloa ruziziensis* gene coexpression network (GCN).

## References

Abdollahi-Arpanahi, R., Gianola, D., & Peñagaricano, F. (2020). Deep learning versus parametric and ensemble methods for genomic prediction of complex phenotypes. Genetics, selection, evolution: GSE, 52(1), 12. 10.1186/s12711-020-00531-z

Agati, G.; Brunetti, C.; Fini, A.; Gori, A.; Guidi, L.; Landi, M.; Sebastiani, F.; Tattini, M. Are Flavonoids Effective Antioxidants in Plants? Twenty Years of Our Investigation. Antioxidants 2020, 9, 1098. 10.3390/antiox9111098

Aguilar-Lopez, J. L., Laboy, R., Jaimes-Miranda, F., Garay, E., DeLuna, A., & Funes, S. (2016). Slm35 links mitochondrial stress response and longevity through TOR signaling pathway. In Aging (Vol. 8, Issue 12, pp. 3255–3271). Impact Journals, LLC. 10.18632/aging.101093

Agullo, G., Gamet-Payrastre, L., Manenti, S., et al. (1997) Relationship between flavonoid structure and inhibition of phosphatidylinositol 3-kinase: a comparison with tyrosine kinase and protein kinase C inhibition. Biochemical Pharmacology, 53, 1649–1657.

Alexa A., Rahnenfuhrer J. (2022). topGO: Enrichment Analysis for Gene Ontology. R package version 2.48.0.

Altschul, S. F., Gish, W., Miller, W., Myers, E. W., & Lipman, D. J. (1990). Basic local alignment search tool. In Journal of Molecular Biology (Vol. 215, Issue 3, pp. 403–410). Elsevier BV. 10.1016/s0022-2836(05)80360-2

Amrine, K. C. H., Blanco-Ulate, B., & Cantu, D. (2015). Discovery of Core Biotic Stress Responsive Genes in Arabidopsis by Weighted Gene Co-Expression Network Analysis. In A. de la Fuente (Ed.), PLOS ONE (Vol. 10, Issue 3, p. e0118731). Public Library of Science (PLoS). 10.1371/journal.pone.0118731

Andrews, S. (2010). FastQC: A Quality Control Tool for High Throughput Sequence Data [Online]. Available online at: http://www.bioinformatics.babraham.ac.uk/projects/fastqc/

Aono, A.H., Costa, E.A., Rody, H.V.S. et al. Machine learning approaches reveal genomic regions associated with sugarcane brown rust resistance. Sci Rep 10, 20057 (2020). 10.1038/s41598-020-77063-5

Aono, A. H., Ferreira, R., Moraes, A., Lara, L., Pimenta, R., Costa, E. A., Pinto, L. R., Landell, M., Santos, M. F., Jank, L., Barrios, S., do Valle, C. B., Chiari, L., Garcia, A., Kuroshu, R. M., Lorena, A. C., Gorjanc, G., & de Souza, A. P. (2022). A joint learning approach for genomic prediction in polyploid grasses. Scientific reports, 12(1), 12499. 10.1038/s41598-022-16417-7

Ashraf, B. H., Jensen, J., Asp, T., & Janss, L. L. (2014). Association studies using family pools of outcrossing crops based on allele-frequency estimates from DNA sequencing. TAG. Theoretical and applied genetics. Theoretische und angewandte Genetik, 127(6), 1331–1341. 10.1007/s00122-014-2300-4

Azodi, C. B., Bolger, E., McCarren, A., Roantree, M., de los Campos, G., & Shiu, S.-H. (2019-A). Benchmarking Parametric and Machine Learning Models for Genomic Prediction of Complex Traits. In G3 Genes|Genomes|Genetics (Vol. 9, Issue 11, pp. 3691–3702). Oxford University Press (OUP). 10.1534/g3.119.400498

Azodi, C. B., Pardo, J., VanBuren, R., de los Campos, G., & Shiu, S.-H. (2019-B). Transcriptome-Based Prediction of Complex Traits in Maize. In The Plant Cell (Vol. 32, Issue 1, pp. 139–151). Oxford University Press (OUP). 10.1105/tpc.19.00332 (B)

Bahri, B.A., Daverdin, G., Xu, X. et al. Natural Variation in Lignin and Pectin Biosynthesis-Related Genes in Switchgrass (Panicum virgatum L.) and Association of SNP Variants with Dry Matter Traits. Bioenerg. Res. 13, 79–99 (2020). 10.1007/s12155-020-10090-2

Baker, C. R., Hanson-Smith, V., & Johnson, A. D. (2013). Following Gene Duplication, Paralog Interference Constrains Transcriptional Circuit Evolution. In Science (Vol. 342, Issue 6154, pp. 104–108). American Association for the Advancement of Science (AAAS). 10.1126/science.1240810

Baskar, V., Venkatesh, R., Ramalingam, S. (2018). Flavonoids (Antioxidants Systems) in Higher Plants and Their Response to Stresses. In: Gupta, D., Palma, J., Corpas, F. (eds) Antioxidants and Antioxidant Enzymes in Higher Plants. Springer, Cham. 10.1007/978-3-319-75088-0_12

Bayer, P. E., Petereit, J., Danilevicz, M. F., Anderson, R., Batley, J., & Edwards, D. (2021). The application of pangenomics and machine learning in genomic selection in plants. In The Plant Genome (Vol. 14, Issue 3). Wiley. 10.1002/tpg2.20112

Barabási A. L., Oltvai Z. N. (2004). Network biology: understanding the cell’s functional organization. Nat. Rev. Genet. 5, 101–113. doi: 10.1038/nrg1272

Bateman, A., Martin, M.-J., Orchard, S., Magrane, M., Agivetova, R., Ahmad, S., Alpi, E., Bowler-Barnett, E. H., Britto, R., Bursteinas, B., Bye-A-Jee, H., Coetzee, R., Cukura, A., Da Silva, A., Denny, P., Dogan, T., Ebenezer, T., Fan, J.,…Teodoro, D. (2020). UniProt: the universal protein knowledgebase in 2021. In Nucleic Acids Research (Vol. 49, Issue D1, pp. D480–D489). Oxford University Press (OUP). 10.1093/nar/gkaa1100

Bélanger, S., Esteves, P., Clermont, I., Jean, M., & Belzile, F. (2016). Genotyping-by-Sequencing on Pooled Samples and its Use in Measuring Segregation Bias during the Course of Androgenesis in Barley. The plant genome, 9(1), 10.3835/plantgenome2014.10.0073. 10.3835/plantgenome2014.10.0073

Bellot, P., de los Campos, G., & Pérez-Enciso, M. (2018). Can Deep Learning Improve Genomic Prediction of Complex Human Traits? In Genetics (Vol. 210, Issue 3, pp. 809–819). Oxford University Press (OUP). 10.1534/genetics.118.301298

Benková, E., Michniewicz, M., Sauer, M., Teichmann, T., Seifertová, D., Jürgens, G., & Friml, J. (2003). Local, Efflux-Dependent Auxin Gradients as a Common Module for Plant Organ Formation. In Cell (Vol. 115, Issue 5, pp. 591–602). Elsevier BV. 10.1016/s0092-8674(03)00924-3

Berro, I., Lado, B., Nalin, R. S., Quincke, M. & Gutiérrez, L. Training population optimization for genomic selection. Plant Genome 12, 190028 (2019).

Bian, Y., & Holland, J. B. (2017). Enhancing genomic prediction with genome-wide association studies in multiparental maize populations. Heredity, 118(6), 585–593. 10.1038/hdy.2017.4

Biazzi E, Nazzicari N, Pecetti L, Brummer EC, Palmonari A, et al. 2017. Genome-wide association mapping and genomic selection for alfalfa (*Medicago sativa*) forage quality traits. PLoS One. 12: e0169234.

Blancaflor, E. B., & Masson, P. H. (2003). Plant Gravitropism. Unraveling the Ups and Downs of a Complex Process. In Plant Physiology (Vol. 133, Issue 4, pp. 1677–1690). Oxford University Press (OUP). 10.1104/pp.103.032169

Blakeslee, J. J., Peer, W. A., & Murphy, A. S. (2005). Auxin transport. In Current Opinion in Plant Biology (Vol. 8, Issue 5, pp. 494–500). Elsevier BV. 10.1016/j.pbi.2005.07.014

Blilou, I., Xu, J., Wildwater, M., Willemsen, V., Paponov, I., Friml, J., Heidstra, R., Aida, M., Palme, K., & Scheres, B. (2005). The PIN auxin efflux facilitator network controls growth and patterning in Arabidopsis roots. In Nature (Vol. 433, Issue 7021, pp. 39–44). Springer Science and Business Media LLC. 10.1038/nature03184

Breiman, L. Bagging predictors. Mach. Learn. 24, 123–140. 10.1007/BF00058655 (2001).

Brown, D.E., Rashotte, A.M., Murphy, A.S. et al. (2001) Flavonoids act as negative regulators of auxin transport in vivo in Arabidopsis. Plant Physiology, 126, 524–535.

Bryant, D. M., Johnson, K., DiTommaso, T., Tickle, T., Couger, M. B., Payzin-Dogru, D., Lee, T. J., Leigh, N. D., Kuo, T.-H., Davis, F. G., Bateman, J., Bryant, S., Guzikowski, A. R., Tsai, S. L., Coyne, S., Ye, W. W., Freeman, R. M., Jr., Peshkin, L., Tabin, C. J.,…Whited, J. L. (2017). A Tissue-Mapped Axolotl De Novo Transcriptome Enables Identification of Limb Regeneration Factors. In Cell Reports (Vol. 18, Issue 3, pp. 762–776). Elsevier BV. 10.1016/j.celrep.2016.12.063

Bolger, A. M., Lohse, M., & Usadel, B. (2014). Trimmomatic: A flexible trimmer for Illumina Sequence Data. Bioinformatics, btu170.

Borin, G. P., Carazzolle, M. F., Dos Santos, R., Riaño-Pachón, D. M., & Oliveira, J. (2018). Gene Co-expression Network Reveals Potential New Genes Related to Sugarcane Bagasse Degradation in *Trichoderma reesei* RUT-30. Frontiers in bioengineering and biotechnology, 6, 151. 10.3389/fbioe.2018.00151

Botté, C. Y., Yamaryo-Botté, Y., Janouškovec, J., Rupasinghe, T., Keeling, P. J., Crellin, P.,…& McFadden, G.I. (2011). Identification of plant-like galactolipids in Chromera velia, a photosynthetic relative of malaria parasites. Journal of Biological Chemistry, 286(34), 29893–29903.

Bout, S., & Vermerris, W. (2003). A candidate-gene approach to clone the sorghum Brown midrib gene encoding caffeic acid O-methyltransferase. In Molecular Genetics and Genomics (Vol. 269, Issue 2, pp. 205–214). Springer Science and Business Media LLC. 10.1007/s00438-003-0824-4

Buell, C. R. (2008). Poaceae Genomes: Going from Unattainable to Becoming a Model Clade for Comparative Plant Genomics. In Plant Physiology (Vol. 149, Issue 1, pp. 111–116). Oxford University Press (OUP). 10.1104/pp.108.128926

Byrne, S., Czaban, A., Studer, B., Panitz, F., Bendixen, C., & Asp, T. (2013). Genome wide allele frequency fingerprints (GWAFFs) of populations via genotyping by sequencing. PloS one, 8(3), e57438. 10.1371/journal.pone.0057438

Cabrera-Bosquet, L., Crossa, J., von Zitzewitz, J., Serret, M. D., & Araus, J. L. (2012). High-throughput phenotyping and genomic selection: the frontiers of crop breeding converge. Journal of integrative plant biology, 54(5), 312–320.

Cai, J., Luo, J., Wang, S. & Yang, S. Feature selection in machine learning: A new perspective. Neurocomputing 300, 70–79 (2018).

Calabrese, G. M., Mesner, L. D., Stains, J. P., Tommasini, S. M., Horowitz, M. C., Rosen, C. J., et al. (2017). Integrating GWAS and co-expression network data identifies bone mineral density genes SPTBN1 and MARK3 and an osteoblast functional module. Cell Syst. 4, 46–59. doi: 10.1016/j.cels.2016.10.014

Cardoso-Silva, C. B., Aono, A. H., Mancini, M. C., Sforça, D. A., da Silva, C. C., Pinto, L. R., Adams, K. L., & de Souza, A. P. (2022). Taxonomically Restricted Genes Are Associated With Responses to Biotic and Abiotic Stresses in Sugarcane (*Saccharum* spp.). Frontiers in plant science, 13, 923069. 10.3389/fpls.2022.923069

Carlson, M. R., Zhang, B., Fang, Z., Mischel, P. S., Horvath, S., & Nelson, S. F. (2006). Gene connectivity, function, and sequence conservation: predictions from modular yeast co-expression networks. In BMC Genomics (Vol. 7, Issue 1). Springer Science and Business Media LLC. 10.1186/1471-2164-7-40

Carpenter, C. D., Kreps, J. A., & Simon, A. E. (1994). Genes Encoding Glycine-Rich Arabidopsis thaliana Proteins with RNA-Binding Motifs Are Influenced by Cold Treatment and an Endogenous Circadian Rhythm. In Plant Physiology (Vol. 104, Issue 3, pp. 1015–1025). Oxford University Press (OUP). 10.1104/pp.104.3.1015

Cericola, F., Lenk, I., Fè, D., Byrne, S., Jensen, C. S., Pedersen, M. G., Asp, T., Jensen, J., & Janss, L. (2018). Optimized Use of Low-Depth Genotyping-by-Sequencing for Genomic Prediction Among Multi-Parental Family Pools and Single Plants in Perennial Ryegrass (Lolium perenne L.). Frontiers in plant science, 9, 369.

Chaffai, R., Elhammadi, M. A., Seybou, T. N., Tekitek, A., Marzouk, B., & El Ferjani, E. (2007). Altered Fatty Acid Profile of Polar Lipids in Maize Seedlings in Response to Excess Copper. In Journal of Agronomy and Crop Science (Vol. 193, Issue 3, pp. 207–217). Wiley. 10.1111/j.1439-037x.2007.00252.x

Chen, T., & Guestrin, C. Xgboost: A scalable tree boosting system. In KDD’16: Proceedings of the 22nd ACM SIGKDD International Conference on Knowledge Discovery and Data Mining. 785–794 (ACM, New York, 2016).

Chen, K., Li, G., Bressan, R. A., Song, C., Zhu, J., & Zhao, Y. (2020). Abscisic acid dynamics, signaling, and functions in plants. In Journal of Integrative Plant Biology (Vol. 62, Issue 1, pp. 25–54). Wiley. 10.1111/jipb.12899

Childs, K. L., Davidson, R. M., and Buell, C. R. (2011). Gene coexpression network analysis as a source of functional annotation for rice genes. PLoS One 6: e22196. doi: 10.1371/journal.pone.0022196

Chung, CW., Hsiao, TH., Huang, CJ. et al. Machine learning approaches for the genomic prediction of rheumatoid arthritis and systemic lupus erythematosus. BioData Mining 14, 52 (2021). 10.1186/s13040-021-00284-5

Colombari-Filho, J. M., M. D. V. Resende, O. P. Morais, A. P. Castro, E. P. Guimarães, J. A. Pereira, M. M. Utumi, and F. Breseghello, 2013: Upland rice breeding in Brazil: a simultaneous genotypic evaluation of stability, adaptability and grain yield. Euphytica 192, 117–129

Costa, C., Schurr, U., Loreto, F., Menesatti, P., & Carpentier, S. (2019). Plant Phenotyping Research Trends, a Science Mapping Approach. Frontiers in plant science, 9, 1933.

Creux, N., & Harmer, S. (2019). Circadian Rhythms in Plants. In Cold Spring Harbor Perspectives in Biology (Vol. 11, Issue 9, p. a034611). Cold Spring Harbor Laboratory. 10.1101/cshperspect.a034611

Cristianini, N. & Shawe-Taylor, J. An Introduction to Support Vector Machines and Other Kernel-Based Learning Methods (Cambridge University Press, 2000).

Crossa, J., Martini, J. W. R., Gianola, D., Pérez-Rodríguez, P., Jarquin, D., Juliana, P., Montesinos-López, O., & Cuevas, J. (2019). Deep Kernel and Deep Learning for Genome-Based Prediction of Single Traits in Multienvironment Breeding Trials. In Frontiers in Genetics (Vol. 10). Frontiers Media SA. 10.3389/fgene.2019.01168

Crossa, J., Pérez-Rodríguez, P., Cuevas, J., Montesinos-López, O., Jarquín, D., de los Campos, G., et al. (2017). Genomic selection in plant breeding: methods, models, and perspectives. Trends Plant Sci. 22, 961–975. doi: 10.1016/j.tplants.2017.08.011

Daetwyler, H. D., Calus, M. P., Pong-Wong, R., de Los Campos, G., & Hickey, J. M. (2013). Genomic prediction in animals and plants: simulation of data, validation, reporting, and benchmarking. Genetics, 193(2), 347–365. 10.1534/genetics.112.147983

Danecek, P., Auton, A., Abecasis, G., Albers, C. A., Banks, E., DePristo, M. A., Handsaker, R. E., Lunter, G., Marth, G. T., Sherry, S. T., McVean, G., & Durbin, R. (2011). The variant call format and VCFtools. In Bioinformatics (Vol. 27, Issue 15, pp. 2156–2158). Oxford University Press (OUP). 10.1093/bioinformatics/btr330

de Bem Oliveira I, Amadeu RR, Ferrão LFV, Muñoz PR. Optimizing whole-genomic prediction for autotetraploid blueberry breeding. Heredity (Edinb). 2020 Dec;125(6):437–448. doi: 10.1038/s41437-020-00357-x. Epub 2020 Oct 19. PMID: 33077896; PMCID: PMC7784927.

de C. Lara, L. A., Santos, M. F., Jank, L., Chiari, L., Vilela, M. de M., Amadeu, R. R., dos Santos, J. P. R., Pereira, G. da S., Zeng, Z.-B., & Garcia, A. A. F. (2019). Genomic Selection with Allele Dosage in Panicum maximum Jacq. In G3 Genes|Genomes|Genetics (Vol. 9, Issue 8, pp. 2463–2475). Oxford University Press (OUP). 10.1534/g3.118.200986 (2020).

De Mendiburu, F., & De Mendiburu, M. F. Package ‘agricolae’. R package version, 1–2

Devos, K. M. (2010). Grass genome organization and evolution. In Current Opinion in Plant Biology (Vol. 13, Issue 2, pp. 139–145). Elsevier BV. 10.1016/j.pbi.2009.12.005

D’haeseleer, P., Liang, S., and Somogyi, R. (2000). Genetic network inference: from co-expression clustering to reverse engineering. Bioinformatics 16, 707–726. doi: 10.1093/bioinformatics/16.8.707

Dhingani RM, Umrania VV, Tomar RS, et al. Introduction to QTL mapping in plants. Ann Plant Sci. 2015;4(04):1072–1079.

Doyle, M.R.; Davis, S.J.; Bastow, R.M.; McWatters, H.G.; Kozma-Bognar, L.; Nagy, F.; Millar, A.J.; Amasino, R.M. The ELF4 gene controls circadian rhythms and flowering time in Arabidopsis thaliana. Nature 2002, 419, 74–77.

Du, H., Zhu, J., Su, H., Huang, M., Wang, H., Ding, S., Zhang, B., Luo, A., Wei, S., Tian, X., & Xu, Y. (2017). Bulked Segregant RNA-seq Reveals Differential Expression and SNPs of Candidate Genes Associated with Waterlogging Tolerance in Maize. Frontiers in plant science, 8, 1022.

Edae, E. A., & Rouse, M. N. (2019). Bulked segregant analysis RNA-seq (BSR-Seq) validated a stem resistance locus in Aegilops umbellulata, a wild relative of wheat. PloS one, 14(9), e0215492.

Edwards, S.M., Buntjer, J.B., Jackson, R. et al. The effects of training population design on genomic prediction accuracy in wheat. Theor Appl Genet 132, 1943–1952 (2019). 10.1007/s00122-019-03327-y

Elkind, Y., Edwards, R., Mavandad, M., Hedrick, S. A., Ribak, O., Dixon, R. A., & Lamb, C. J. (1990). Abnormal plant development and down-regulation of phenylpropanoid biosynthesis in transgenic tobacco containing a heterologous phenylalanine ammonia-lyase gene. In Proceedings of the National Academy of Sciences (Vol. 87, Issue 22, pp. 9057–9061). Proceedings of the National Academy of Sciences. 10.1073/pnas.87.22.9057

Elshire, R. J., Glaubitz, J. C., Sun, Q., Poland, J. A., Kawamoto, K., Buckler, E. S., et al. (2011). A robust, simple genotyping-by-sequencing (GBS) approach for high diversity species. PLoS One 6:e19379. doi: 10.1371/journal.pone.0019379

Eves-van den Akker, S. (2021). Plant–nematode interactions. In Current Opinion in Plant Biology (Vol. 62, p. 102035). Elsevier BV. 10.1016/j.pbi.2021.102035

Farrow, S. C., & Facchini, P. J. (2014). Functional diversity of 2-oxoglutarate/Fe(II)-dependent dioxygenases in plant metabolism. In Frontiers in Plant Science (Vol. 5). Frontiers Media SA. 10.3389/fpls.2014.00524

Fè, D., Cericola, F., Byrne, S. et al. Genomic dissection and prediction of heading date in perennial ryegrass. BMC Genomics 16, 921 (2015). 10.1186/s12864-015-2163-3

Fernández-Marcos, M., Sanz, L., Lewis, D.R., et al. (2013) Control of auxin transport by reactive oxygen and nitrogen species, in Polar Auxin Transport, Signaling and Communication in Plants, vol. 17 (eds R. Chen and F. Baluska), Springer-Verlag, Berlin, pp. 103–117

Ferrão, L., Amadeu, R. R., Benevenuto, J., de Bem Oliveira, I., & Munoz, P. R. (2021). Genomic Selection in an Outcrossing Autotetraploid Fruit Crop: Lessons From Blueberry Breeding. Frontiers in plant science, 12, 676326. 10.3389/fpls.2021.676326

Ferreira, R. C. U., Cançado, L. J., Do Valle, C. B., Chiari, L., and de Souza, A. P. (2016). Microsatellite loci for Urochloa decumbens (Stapf) R.D. Webster and cross-amplification in other Urochloa species. BMC. Res. Notes 9:152. doi: 10.1186/s13104-016-1967-9

Ferreira R. C. U, da Costa Lima Moraes A, Chiari L, Simeão RM, Vigna BBZ, de Souza AP. An Overview of the Genetics and Genomics of the Urochloa Species Most Commonly Used in Pastures. Front Plant Sci. 2021 Dec 13; 12:770461. doi: 10.3389/fpls.2021.770461. PMID: 34966402; PMCID: PMC8710810.

Figueiredo, U. J. de, Nunes, J. A. R., & Valle, C. B. do. (2012). Estimation of genetic parameters and selection of Brachiaria humidicola progenies using a selection index. In Crop Breeding and Applied Biotechnology (Vol. 12, Issue 4, pp. 237–244). FapUNIFESP (SciELO). 10.1590/s1984-70332012000400002

Force, A., Lynch, M., Pickett, F. B., Amores, A., Yan, Y., & Postlethwait, J. (1999). Preservation of Duplicate Genes by Complementary, Degenerative Mutations. In Genetics (Vol. 151, Issue 4, pp. 1531–1545). Oxford University Press (OUP). 10.1093/genetics/151.4.1531

Francisco, F. R., Aono, A. H., da Silva, C. C., Gonçalves, P. S., Scaloppi Junior, E. J., Le Guen, V., Fritsche-Neto, R., Souza, L. M., & de Souza, A. P. (2021). Unravelling Rubber Tree Growth by Integrating GWAS and Biological Network-Based Approaches. Frontiers in plant science, 12, 768589. 10.3389/fpls.2021.768589

Franke, R., Hemm, M. R., Denault, J. W., Ruegger, M. O., Humphreys, J. M., & Chapple, C. (2002). Changes in secondary metabolism and deposition of an unusual lignin in the ref8 mutant of Arabidopsis. In The Plant Journal (Vol. 30, Issue 1, pp. 47–59). Wiley. 10.1046/j.1365-313x.2002.01267.x

Freeling, M., & Thomas, B. C. (2006). Gene-balanced duplications, like tetraploidy, provide predictable drive to increase morphological complexity. In Genome Research (Vol. 16, Issue 7, pp. 805–814). Cold Spring Harbor Laboratory. 10.1101/gr.3681406

Freund, Y. & Schapire, R. E. A decision-theoretic generalization of on-line learning and an application to boosting. J. Comput. Syst. Sci. 55, 119–139. 10.1006/jcss.1997.1504 (1997).

Friml J, Vieten A, Sauer M, Weijers D, Schwarz H, et al. 2003. Efflux-dependent auxin gradients establish the apical-basal axis of Arabidopsis. Nature 426:147–53

Fujikura, U., Horiguchi, G., Ponce, M. R., Micol, J. L., & Tsukaya, H. (2009). Coordination of cell proliferation and cell expansion mediated by ribosome-related processes in the leaves of Arabidopsis thaliana. The Plant Journal, 59(3), 499– 508. 10.1111/j.1365-313X.2009.03886.x

Futschik, A., & Schlötterer, C. (2010). The Next Generation of Molecular Markers From Massively Parallel Sequencing of Pooled DNA Samples. In Genetics (Vol. 186, Issue 1, pp. 207–218). Oxford University Press (OUP). 10.1534/genetics.110.114397

Gaut, B. S. (2002). Evolutionary dynamics of grass genomes. In New Phytologist (Vol. 154, Issue 1, pp. 15–28). Wiley. 10.1046/j.1469-8137.2002.00352.x

Geurts, P., Ernst, D. & Wehenkel, L. Extremely randomized trees. Mach. Learn. 63, 3–42. 10.1007/s10994-006-6226-1 (2006).

Glaubitz, J. C., Casstevens, T. M., Lu, F., Harriman, J., Elshire, R. J., Sun, Q., et al. (2014). TASSEL-GBS: a high capacity genotyping by sequencing analysis pipeline. PloS One 9, e90346. doi: 10.1371/journal.pone.0090346

Goddard, M. E., Kemper, K. E., MacLeod, I. M., Chamberlain, A. J., & Hayes, B. J. (2016). Genetics of complex traits: prediction of phenotype, identification of causal polymorphisms and genetic architecture. Proceedings. Biological sciences, 283(1835), 20160569. 10.1098/rspb.2016.0569

Gorelova, V.; Ambach, L.; Rébeillé, F.; Stove, C.; Van Der Straeten, D. Folates in plants: Research advances and progress in crop biofortification. Front. Chem. 2017, 5, 21.

Grabherr, M. G., Haas, B. J., Yassour, M., Levin, J. Z., Thompson, D. A., Amit, I., Adiconis, X., Fan, L., Raychowdhury, R., Zeng, Q., Chen, Z., Mauceli, E., Hacohen, N., Gnirke, A., Rhind, N., di Palma, F., Birren, B. W., Nusbaum, C., Lindblad-Toh, K.,…Regev, A. (2011). Full-length transcriptome assembly from RNA-Seq data without a reference genome. Nature Biotechnology, 29(7), 644–652. 10.1038/nbt.1883

Granato, I., Cuevas, J., Luna-Vázquez, F., Crossa, J., Montesinos-López, O., Burgueño, J., & Fritsche-Neto, R. (2018). BGGE: A New Package for Genomic-Enabled Prediction Incorporating Genotype × Environment Interaction Models. In G3 Genes|Genomes|Genetics (Vol. 8, Issue 9, pp. 3039–3047). Oxford University Press (OUP). 10.1534/g3.118.200435

Genuth, N. R., & Barna, M. (2018). The discovery of ribosome heterogeneity and its implications for gene regulation and organismal life. Molecular Cell, 71(3), 364–374. 10.1016/j.molcel.2018.07.018

Gigon, A., Matos, A.-R., Laffray, D., Zuily-Fodil, Y., & Pham-Tthi, A.-T. (2004). Effect of Drought Stress on Lipid Metabolism in the Leaves of Arabidopsis thaliana (Ecotype Columbia). In Annals of Botany (Vol. 94, Issue 3, pp. 345–351). Oxford University Press (OUP). 10.1093/aob/mch150

Gilbert, W. V. (2011). Functional specialization of ribosomes? Trends in Biochemical Sciences, 36(3), 127– 132. 10.1016/j.tibs.2010.12.002

Grieneisen VA, Xu J, Maree AF, Hogeweg P, Scheres B. 2007. Auxin transport is sufficient to generate a maximum and gradient guiding root growth. Nature 449:1008–13

Grinberg, N. F., Lovatt, A., Hegarty, M., Lovatt, A., Skøt, K. P., Kelly, R., Blackmore, T., Thorogood, D., King, R. D., Armstead, I., Powell, W., & Skøt, L. (2016). Implementation of Genomic Prediction in Lolium perenne (L.) Breeding Populations. In Frontiers in Plant Science (Vol. 7). Frontiers Media SA. 10.3389/fpls.2016.00133

Grunewald, W., De Smet, I., Lewis, D.R. et al. (2012) Transcription factor WRKY23 assists auxin distribution patterns during Arabidopsis root development through local control on flavonol biosynthesis. Proceedings of the National Academy of Sciences of the United States of America, 109, 1554–1559.

Gu Z, Gu L, Eils R, Schlesner M, Brors B. circlize Implements and enhances circular visualization in R. Bioinformatics. 2014 Oct;30(19):2811–2. doi: 10.1093/bioinformatics/btu393. Epub 2014 Jun 14. PMID: 24930139.

Guo X, Cericola F, Fè D, Pedersen MG, Lenk I, et al. 2018. Genomic prediction in tetraploid ryegrass using allele frequencies based on genotyping by sequencing. Front Plant Sci. 9:1165.

Guo, Z., Zhou, S., Wang, S., Li, W. X., Du, H., & Xu, Y. (2021). Identification of major QTL for waterlogging tolerance in maize using genome-wide association study and bulked sample analysis. Journal of applied genetics, 62(3), 405–418.

Haile TA, Walkowiak S, N’Diaye A, Clarke JM, Hucl PJ, Cuthbert RD, Knox RE, Pozniak CJ. Genomic prediction of agronomic traits in wheat using different models and cross-validation designs. Theor Appl Genet. 2021 Jan;134(1):381–398. doi: 10.1007/s00122-020-03703-z. Epub 2020 Nov 1. PMID: 33135095.

Hanley, S. J., Pellny, T. K., de Vega, J. J., Castiblanco, V., Arango, J., Eastmond, P. J., Heslop-Harrison, J. S. P., & Mitchell, R. A. C. (2021). Allele mining in diverse accessions of tropical grasses to improve forage quality and reduce environmental impact. Annals of Botany, 128(5), 627–637. 10.1093/aob/mcab101

Heer, K., Behringer, D., Piermattei, A., Bässler, C., Brandl, R., Fady, B., Jehl, H., Liepelt, S., Lorch, S., Piotti, A., Vendramin, G. G., Weller, M., Ziegenhagen, B., Büntgen, U., & Opgenoorth, L. (2018). Linking dendroecology and association genetics in natural populations: Stress responses archived in tree rings associate with SNP genotypes in silver fir (Abies albaMill.). In Molecular Ecology (Vol. 27, Issue 6, pp. 1428–1438). Wiley. 10.1111/mec.14538

Horiguchi, G., Mollá-Morales, A., Pérez-Pérez, J. M., Kojima, K., Robles, P., Ponce, M. R., Micol, J. L., & Tsukaya, H. (2011). Differential contributions of ribosomal protein genes to Arabidopsis thaliana leaf development. The Plant Journal, 65(5), 724– 736. 10.1111/j.1365-313X.2010.04457.x

Hou, G., Ablett, G. R., Pauls, K. P., & Rajcan, I. (2006). Environmental effects on fatty acid levels in soybean seed oil. In Journal of the American Oil Chemists’ Society (Vol. 83, Issue 9, pp. 759–763). Wiley. 10.1007/s11746-006-5011-4

Hou, Q., Ufer, G., & Bartels, D. (2016). Lipid signalling in plant responses to abiotic stress. In Plant, Cell Environment (Vol. 39, Issue 5, pp. 1029–1048). Wiley. 10.1111/pce.12666

Iba, K. (2002). Acclimative response to temperature stress in higher plants: Approaches of Gene Engineering for Temperature Tolerance. In Annual Review of Plant Biology (Vol. 53, Issue 1, pp. 225–245). Annual Reviews. 10.1146/annurev.arplant.53.100201.160729

Islam, M. S., McCord, P. H., Olatoye, M. O., Qin, L., Sood, S., Lipka, A. E., & Todd, J. R. (2021). Experimental evaluation of genomic selection prediction for rust resistance in sugarcane. In The Plant Genome (Vol. 14, Issue 3). Wiley. 10.1002/tpg2.20148

Ito, T., Kim, G. T., & Shinozaki, K. (2000). Disruption of an Arabidopsis cytoplasmic ribosomal protein S13-homologous gene by transposon-mediated mutagenesis causes aberrant growth and development. The Plant Journal, 22(3), 257–264. 10.1046/j.1365-313x.2000.00728.x

Jacobs, M. and Rubery, P.H. (1988) Naturally-occurring auxin transport regulators. Science, 241, 346–349

Jank L., Barrios S. C., do Valle C. B., Simeão R. M., Alves G. F. (2014). The value of improved pastures to Brazilian beef production. Crop Pasture Sci. 65, 1132–1137. doi: 10.1071/CP13319

Jia, C., Zhao, F., Wang, X., Han, J., Zhao, H., Liu, G., & Wang, Z. (2018). Genomic Prediction for 25 Agronomic and Quality Traits in Alfalfa (Medicago sativa). Frontiers in plant science, 9, 1220.

Jeong, S., Kim, J. Y., & Kim, N. (2020). GMStool: GWAS-based marker selection tool for genomic prediction from genomic data. Scientific reports, 10(1), 19653. 10.1038/s41598-020-76759-y

Juliana P, He X, Marza F, Islam R, Anwar B, Poland J, Shrestha S, Singh GP, Chawade A, Joshi AK, Singh RP and Singh PK (2022) Genomic Selection for Wheat Blast in a Diversity Panel, Breeding Panel and Full-Sibs Panel. Front. Plant Sci. 12:745379. doi: 10.3389/fpls.2021.745379

Jiao, Y., Zhao, H., Ren, L., Song, W., Zeng, B., Guo, J., Wang, B., Liu, Z., Chen, J., Li, W., Zhang, M., Xie, S., & Lai, J. (2012). Genome-wide genetic changes during modern breeding of maize. Nature genetics, 44(7), 812–815.

Jones, C., De Vega, J., Worthington, M., Thomas, A., Gasior, D., Harper, J., et al. (2021). A comparison of differential gene expression in response to the onset of water stress between three hybrid Brachiaria genotypes. Front. Plant Sci. 12:637956. doi: 10.3389/fpls.2021.637956

Jones, L., Ennos, A. R., & Turner, S. R. (2001). Cloning and characterization of irregular xylem4 (irx4): a severely lignin-deficient mutant of Arabidopsis. In The Plant Journal (Vol. 26, Issue 2, pp. 205–216). Wiley. 10.1046/j.1365-313x.2001.01021.x

Jouhet, J., Maréchal, E., & Block, M. A. (2007). Glycerolipid transfer for the building of membranes in plant cells. Progress in lipid research, 46(1), 37–55.

Kachroo, A., Lapchyk, L., Fukushige, H., Hildebrand, D., Klessig, D., & Kachroo, P. (2003). Plastidial Fatty Acid Signaling Modulates Salicylic Acid– and Jasmonic Acid–Mediated Defense Pathways in the Arabidopsis ssi2 Mutant. In The Plant Cell (Vol. 15, Issue 12, pp. 2952–2965). Oxford University Press (OUP). 10.1105/tpc.017301

Karthika, V., Babitha, K.C., Kiranmai, K. et al. Involvement of DNA mismatch repair systems to create genetic diversity in plants for speed breeding programs. Plant Physiol. Rep. 25, 185–199 (2020). 10.1007/s40502-020-00521-9

Kim, J., Kim, H.-S., Choi, S.-H., Jang, J.-Y., Jeong, M.-J., & Lee, S. (2017). The Importance of the Circadian Clock in Regulating Plant Metabolism. In International Journal of Molecular Sciences (Vol. 18, Issue 12, p. 2680). MDPI AG. 10.3390/ijms18122680

Kim, M. K., & Kim, W. T. (2018). Telomere Structure, Function, and Maintenance in Plants. In Journal of Plant Biology (Vol. 61, Issue 3, pp. 131–136). Springer Science and Business Media LLC. 10.1007/s12374-018-0082-y

Kobayashi, K., Kondo, M., Fukuda, H., Nishimura, M., & Ohta, H. (2007). Galactolipid synthesis in chloroplast inner envelope is essential for proper thylakoid biogenesis, photosynthesis, and embryogenesis. In Proceedings of the National Academy of Sciences (Vol. 104, Issue 43, pp. 17216–17221). Proceedings of the National Academy of Sciences. 10.1073/pnas.0704680104

Kombrink, A., Sánchez-Vallet, A., & Thomma, B. P. H. J. (2011). The role of chitin detection in plant–pathogen interactions. In Microbes and Infection (Vol. 13, Issues 14–15, pp. 1168–1176). Elsevier BV. 10.1016/j.micinf.2011.07.010

Korte, A., Farlow, A. The advantages and limitations of trait analysis with GWAS: a review. Plant Methods 9, 29 (2013). 10.1186/1746-4811-9-29

Lauvergeat, V., Lacomme, C., Lacombe, E., Lasserre, E., Roby, D., & Grima-Pettenati, J. (2001). Two cinnamoyl-CoA reductase (CCR) genes from Arabidopsis thaliana are differentially expressed during development and in response to infection with pathogenic bacteria. In Phytochemistry (Vol. 57, Issue 7, pp. 1187–1195). Elsevier BV. 10.1016/s0031-9422(01)00053-x

Li, B., Zhang, N., Wang, Y.-G., George, A. W., Reverter, A., & Li, Y. (2018). Genomic Prediction of Breeding Values Using a Subset of SNPs Identified by Three Machine Learning Methods. In Frontiers in Genetics (Vol. 9). Frontiers Media SA. 10.3389/fgene.2018.00237

Li, X., Yang, Y., Yao, J. et al. FLEXIBLE CULM 1 encoding a cinnamyl-alcohol dehydrogenase controls culm mechanical strength in rice. Plant Mol Biol 69, 685–697 (2009). 10.1007/s11103-008-9448-8

Liang, M., Miao, J., Wang, X., Chang, T., An, B., Duan, X., Xu, L., Gao, X., Zhang, L., Li, J., & Gao, H. (2020). Application of ensemble learning to genomic selection in chinese simmental beef cattle. In Journal of Animal Breeding and Genetics (Vol. 138, Issue 3, pp. 291–299). Wiley. 10.1111/jbg.12514

Liang, M., Chang, T., An, B., Duan, X., Du, L., Wang, X., Miao, J., Xu, L., Gao, X., Zhang, L., Li, J., & Gao, H. (2021). A Stacking Ensemble Learning Framework for Genomic Prediction. In Frontiers in Genetics (Vol. 12). Frontiers Media SA. 10.3389/fgene.2021.600040

Lipka, A. E., Lu, F., Cherney, J. H., Buckler, E. S., Casler, M. D., & Costich, D. E. (2014). Accelerating the Switchgrass (Panicum virgatum L.) Breeding Cycle Using Genomic Selection Approaches. In D. D. Fang (Ed.), PLoS ONE (Vol. 9, Issue 11, p. e112227). Public Library of Science (PLoS). 10.1371/journal.pone.0112227

Liu, S., Feuerstein, U., Luesink, W., Schulze, S., Asp, T., Studer, B., Becker, H. C., & Dehmer, K. J. (2018). DArT, SNP, and SSR analyses of genetic diversity in Lolium perenne L. using bulk sampling. BMC genetics, 19(1), 10.

Liu, C., Ma, Y., Zhao, J., Nussinov, R., Zhang, Y. C., Cheng, F., et al. (2020). Computational network biology: data, models, and applications. Phys. Rep. 846, 1–66. doi: 10.1016/j.physrep.2019.12.004

Lorenz, A. J., & Smith, K. P. (2015). Adding Genetically Distant Individuals to Training Populations Reduces Genomic Prediction Accuracy in Barley. In Crop Science (Vol. 55, Issue 6, pp. 2657–2667). Wiley. 10.2135/cropsci2014.12.0827

Louis Lello, Steven G Avery, Laurent Tellier, Ana I Vazquez, Gustavo de los Campos, Stephen D H Hsu, Accurate Genomic Prediction of Human Height, Genetics, Volume 210, Issue 2, 1 October 2018, Pages 477–497, 10.1534/genetics.118.301267

Luo, Z., Yu, Y., Xiang, J. & Li, F. Genomic selection using a subset of snps identified by genome-wide association analysis for disease resistance traits in aquaculture species. Aquaculture 539, 736620 (2021).

Ma, W., Qiu, Z., Song, J. et al. A deep convolutional neural network approach for predicting phenotypes from genotypes. Planta 248, 1307–1318 (2018). 10.1007/s00425-018-2976-9

Macovei, A., Vaid, N., Tula, S., & Tuteja, N. (2012). A new DEAD-box helicase ATP-binding protein (OsABP) from rice is responsive to abiotic stress. In Plant Signaling & Behavior (Vol. 7, Issue 9, pp. 1138–1143). Informa UK Limited. 10.4161/psb.21343

Maksymiec, W. (2007). Signaling responses in plants to heavy metal stress. In Acta Physiologiae Plantarum (Vol. 29, Issue 3, pp. 177–187). Springer Science and Business Media LLC. 10.1007/s11738-007-0036-3

Mateescu, R. G., Garrick, D. J., & Reecy, J. M. (2017). Network Analysis Reveals Putative Genes Affecting Meat Quality in Angus Cattle. Frontiers in genetics, 8, 171. 10.3389/fgene.2017.00171

Martins, F. B., Moraes, A. C. L., Aono, A. H., Ferreira, R. C. U., Chiari, L., Simeão, R. M., Barrios, S. C. L., Santos, M. F., Jank, L., do Valle, C. B., Vigna, B. B. Z., & de Souza, A. P. (2021). A Semi-Automated SNP-Based Approach for Contaminant Identification in Biparental Polyploid Populations of Tropical Forage Grasses. In Frontiers in Plant Science (Vol. 12). Frontiers Media SA. 10.3389/fpls.2021.737919

Matias, F. I., Alves, F. C., Meireles, K. G. X., Barrios, S. C. L., do Valle, C. B., Endelman, J. B., & Fritsche-Neto, R. (2019a). On the accuracy of genomic prediction models considering multi-trait and allele dosage in Urochloa spp. interspecific tetraploid hybrids. Molecular Breeding, 39(7), 1–16.

Matias, F. I., Vidotti, M. S., Meireles, K. G. X., Barrios, S. C. L., do Valle, C. B., Carley, C. A. S., et al. (2019b). Association mapping considering allele dosage: an example of forage traits in an interspecific segmental allotetraploid Urochloa spp. panel. Crop Sci. 59, 2062–2076. doi: 10.2135/cropsci2019.03.0185

Medina, C. A., Kaur, H., Ray, I., & Yu, L.-X. (2021). Strategies to Increase Prediction Accuracy in Genomic Selection of Complex Traits in Alfalfa (Medicago sativa L.). In Cells (Vol. 10, Issue 12, p. 3372). MDPI AG. 10.3390/cells10123372

Meuwissen THE, Hayes BJ, Goddard ME. 2001. Prediction of total genetic value using genome-wide dense marker maps. Genetics. 157:1819–1829.

Miao, J. & Niu, L. A survey on feature selection. Procedia Comput. Sci. 91, 919–926 (2016).

Mikami, K., & Murata, N. (2003). Membrane fluidity and the perception of environmental signals in cyanobacteria and plants. In Progress in Lipid Research (Vol. 42, Issue 6, pp. 527–543). Elsevier BV. 10.1016/s0163-7827(03)00036-5

Millar AJ. 2016. The intracellular dynamics of circadian clocks reach for the light of ecology and evolution. Annu Rev Plant Biol 67: 595–618.doi:10.1146/annurev-arplant-043014-115619

Montesinos-López, O.A., Montesinos-López, A., Pérez-Rodríguez, P. et al. A review of deep learning applications for genomic selection. BMC Genomics 22, 19 (2021). 10.1186/s12864-020-07319-x

Mohan M, Nair S, Bhagwat A, et al. Genome mapping, molecular markers and marker-assisted selection in crop plants. Mol Breed. 1997;3(2):87–103.

Montesinos-López, O. A., Montesinos-López, A., Tuberosa, R., Maccaferri, M., Sciara, G., Ammar, K., & Crossa, J. (2019). Multi-Trait, Multi-Environment Genomic Prediction of Durum Wheat With Genomic Best Linear Unbiased Predictor and Deep Learning Methods. In Frontiers in Plant Science (Vol. 10). Frontiers Media SA. 10.3389/fpls.2019.01311

Mosè Manni, Matthew R Berkeley, Mathieu Seppey, Felipe A Simão, Evgeny M Zdobnov, BUSCO Update: Novel and Streamlined Workflows along with Broader and Deeper Phylogenetic Coverage for Scoring of Eukaryotic, Prokaryotic, and Viral Genomes, Molecular Biology and Evolution, Volume 38, Issue 10, October 2021, Pages 4647–4654

Murad Leite Andrade, M. H., Acharya, J. P., Benevenuto, J., de Bem Oliveira, I., Lopez, Y., Munoz, P., Resende, M. F. R., Jr., & Rios, E. F. (2022). Genomic prediction for canopy height and dry matter yield in alfalfa using family bulks. In The Plant Genome. Wiley. 10.1002/tpg2.20235

Mutwil, M., Usadel, B., Schultte, M., Loraine, A., Ebenholh, O., & Persson, S. (2009). Assembly of an Interactive Correlation Network for the Arabidopsis Genome Using a Novel Heuristic Clustering Algorithm. In Plant Physiology (Vol. 152, Issue 1, pp. 29–43). Oxford University Press (OUP). 10.1104/pp.109.145318

Nandi, A., Moeder, W., Kachroo, P., Klessig, D. F., & Shah, J. (2005). Arabidopsis ssi2-Conferred Susceptibility to Botrytis cinerea Is Dependent on EDS5 and PAD4. In Molecular Plant-Microbe Interactions® (Vol. 18, Issue 4, pp. 363–370). Scientific Societies. 10.1094/mpmi-18-0363

Norris, K., Hopes, T., & Aspden, J. L. (2021). Ribosome heterogeneity and specialization in development. In WIREs RNA (Vol. 12, Issue 4). Wiley. 10.1002/wrna.1644

Ohno S (1970) Evolution by Gene Duplication. Springer-Verlag, New York

Oka, T., Nemoto, T., & Jigami, Y. (2007). Functional Analysis of Arabidopsis thaliana RHM2/MUM4, a Multidomain Protein Involved in UDP-D-glucose to UDP-L-rhamnose Conversion. In Journal of Biological Chemistry (Vol. 282, Issue 8, pp. 5389–5403). Elsevier BV. 10.1074/jbc.m610196200

Oliver, S. (2000) Guilt-by-association goes global. Nature, 403, 601–602. 10.1038/35001165

Panchy, N., Lehti-Shiu, M., & Shiu, S.-H. (2016). Evolution of Gene Duplication in Plants. In Plant Physiology (Vol. 171, Issue 4, pp. 2294–2316). Oxford University Press (OUP). 10.1104/pp.16.00523

Parker Gaddis, K. L., Null, D. J., & Cole, J. B. (2016). Explorations in genome-wide association studies and network analyses with dairy cattle fertility traits. Journal of dairy science, 99(8), 6420–6435. 10.3168/jds.2015-10444

Patro, R., Duggal, G., Love, M. I., Irizarry, R. A., & Kingsford, C. (2017). Salmon provides fast and bias-aware quantification of transcript expression. Nature methods, 14(4), 417–419.

Pedregosa, F. et al. Scikit-learn: Machine learning in Python. J. Mach. Learn. Res. 12, 2825–2830 (2011).

Peer, W.A. and Murphy, A.S. (2007) Flavonoids and auxin transport: modulators or regulators? Trends in Plant Science, 12, 556–563.

Peer, W.A., Bandyopadhyay, A., Blakeslee, J.J. et al. (2004) Variation in expression and protein localization of the PIN family of auxin efflux facilitator proteins in flavonoid mutants with altered auxin transport in Arabidopsis thaliana. The Plant Cell, 16, 1898–1911..

Pereira, G. S., Garcia, A. A. F., Margarido, G. R. A. (2018a). A fully automated pipeline for quantitative genotype calling from next generation sequencing data in autopolyploids. BMC Bioinform. 19, 398. doi: 10.1186/s12859-018-2433-6

Pereira J. F., Azevedo A. L. S., Pessoa-Filho M., Romanel E. A. D. C., Pereira A. V., Vigna B. B. Z., et al. (2018b). Research priorities for next-generation breeding of tropical forages in Brazil. Crop Breed. Appl. Biotechnol. 18 314–319. 10.1590/1984-70332018v18n3n46

Perez, P., and de los Campos, G., 2014 Genome-Wide Regression and Prediction with the BGLR Statistical Package. Genetics 198 (2): 483–495.

Petrasch S, Mesquida-Pesci SD, Pincot DDA, Feldmann MJ, López CM, Famula R, Hardigan MA, Cole GS, Knapp SJ, Blanco-Ulate B. Genomic prediction of strawberry resistance to postharvest fruit decay caused by the fungal pathogen Botrytis cinerea. G3 (Bethesda). 2022 Jan 4;12(1): jkab378. doi: 10.1093/g3journal/jkab378. PMID: 34791166; PMCID: PMC8728004.

Pessoa-Filho, M., Sobrinho, F. S., Fragoso, R. R., Silva Junior, O. B., and Ferreira, M. E. (2019). “A Phased Diploid Genome Assembly for the Forage Grass Urochloa Ruziziensis Based on Single-Molecule Real-Time Sequencing.” in International Plant and Animal Genome Conference XXVII, 2019, San Diego. Available at: https://www.embrapa.br/en/busca-de-publicacoes/-/publicacao/1107378/a-phased-diploid-genome-assembly-for-the-forage-grass-urochloa-ruziziensis-based-on-single-molecule-real-time-sequencing.

Piles, M., Bergsma, R., Gianola, D., Gilbert, H., & Tusell, L. (2021). Feature Selection Stability and Accuracy of Prediction Models for Genomic Prediction of Residual Feed Intake in Pigs Using Machine Learning. In Frontiers in Genetics (Vol. 12). Frontiers Media SA. 10.3389/fgene.2021.611506

Pimenta, R.J.G., Aono, A.H., Burbano, R.C.V. et al. Genome-wide approaches for the identification of markers and genes associated with sugarcane yellow leaf virus resistance. Sci Rep 11, 15730 (2021). 10.1038/s41598-021-95116-1

Pimenta, R. J. G., Aono, A. H., Burbano, R. C. V., da Silva, M. F., dos Anjos, I. A., de Andrade Landell, M. G., Gonçalves, M. C., Pinto, L. R., & de Souza, A. P. (2022). Multiomic investigation of sugarcane mosaic virus resistance in sugarcane. Cold Spring Harbor Laboratory. 10.1101/2022.08.18.504288

Pincot, D. D. A., Hardigan, M. A., Cole, G. S., Famula, R. A., Henry, P. M., Gordon, T. R., et al. (2020). Accuracy of genomic selection and long−term genetic gain for resistance to *Verticillium* wilt in strawberry. Plant Genome 13: e20054. doi: 10.1002/tpg2.20054

Piquemal, J., Lapierre, C., Myton, K., O’connell, A., Schuch, W., Grima-pettenati, J., & Boudet, A.-M. (2002). Down-regulation of Cinnamoyl-CoA Reductase induces significant changes of lignin profiles in transgenic tobacco plants. In The Plant Journal (Vol. 13, Issue 1, pp. 71–83). Wiley. 10.1046/j.1365-313x.1998.00014.x

Poland, J. A., Brown, P. J., Sorrells, M. E., & Jannink, J. L. (2012). Development of high-density genetic maps for barley and wheat using a novel two-enzyme genotyping-by-sequencing approach. PloS one, 7(2), e32253.

Popescu, M. C., Balas, V., Perescu-Popescu, L. & Mastorakis, N. Multilayer perceptron and neural networks. WSEAS Trans. Circuits Syst. 8, 579–588 (2009).

R Core Team (2021). R: A language and environment for statistical computing. R Foundation for Statistical Computing, Vienna, Austria. URL https://www.R-project.org/.

Rao, X., & Dixon, R. A. (2019). Co-expression networks for plant biology: why and how. Acta biochimica et biophysica Sinica, 51(10), 981–988. 10.1093/abbs/gmz080

Resende, M. D. V., 2002: Software Selegen – REML/BLUP. Embrapa Florestas, Colombo-Brazil.

Reverter, A., & Chan, E. K. F. (2008). Combining partial correlation and an information theory approach to the reversed engineering of gene co-expression networks. In Bioinformatics (Vol. 24, Issue 21, pp. 2491–2497). Oxford University Press (OUP). 10.1093/bioinformatics/btn482

Rios, E. F., Andrade, M. H. M. L., Resende, M. F. R., Jr, Kirst, M., de Resende, M. D. V., de Almeida Filho, J. E., Gezan, S. A., & Munoz, P. (2021). Genomic prediction in family bulks using different traits and cross-validations in pine. In A. E. Lipka (Ed.), G3 Genes|Genomes|Genetics (Vol. 11, Issue 9). Oxford University Press (OUP). 10.1093/g3journal/jkab249

Rosolen, R. R., Aono, A. H., Almeida, D. A., Ferreira Filho, J. A., Horta, M., & De Souza, A. P. (2022). Network Analysis Reveals Different Cellulose Degradation Strategies Across *Trichoderma harzianum* Strains Associated With XYR1 and CRE1. Frontiers in genetics, 13, 807243. 10.3389/fgene.2022.807243

Routaboul, J.-M., Fischer, S. F., & Browse, J. (2000). Trienoic Fatty Acids Are Required to Maintain Chloroplast Function at Low Temperatures. In Plant Physiology (Vol. 124, Issue 4, pp. 1697–1705). Oxford University Press (OUP). 10.1104/pp.124.4.1697

Saballos, A., Sattler, S. E., Sanchez, E., Foster, T. P., Xin, Z., Kang, C., Pedersen, J. F., & Vermerris, W. (2012). Brown midrib2 (Bmr2) encodes the major 4-coumarate:coenzyme A ligase involved in lignin biosynthesis in sorghum (Sorghum bicolor (L.) Moench). In The Plant Journal (Vol. 70, Issue 5, pp. 818–830). Wiley. 10.1111/j.1365-313x.2012.04933.x

Salgado, L. R., Lima, R., Santos, B. F. D., Shirakawa, K. T., Vilela, M. D. A., Almeida, N. F., et al. (2017). De novo RNA sequencing and analysis of the transcriptome of signalgrass (Urochloa decumbens) roots exposed to aluminum. Plant Growth Regul. 83, 157–170. doi: 10.1007/s10725-017-0291-2

Sánchez-Vallet, A., Mesters, J. R., & Thomma, B. P. H. J. (2015). The battle for chitin recognition in plant-microbe interactions. In FEMS Microbiology Reviews4 (Vol. 39, Issue 2, pp. 171–183). Oxford University Press (OUP). 10.1093/femsre/fuu003

Sandhu, K., Aoun, M., Morris, C., & Carter, A. (2021). Genomic Selection for End-Use Quality and Processing Traits in Soft White Winter Wheat Breeding Program with Machine and Deep Learning Models. In Biology (Vol. 10, Issue 7, p. 689). MDPI AG. 10.3390/biology10070689

Santelia, D., Henrichs, S., Vincenzetti, V. et al. (2008) Flavonoids redirect PIN-mediated polar auxin fluxes during root gravitropic responses. Journal of Biological Chemistry, 283, 31218–31226.

Schaefer, R. J., Michno, J. M., Jeffers, J., Hoekenga, O., Dilkes, B., Baxter, I., et al. (2018). Integrating coexpression networks with GWAS to prioritize causal genes in maize. Plant Cell 30, 2922–2942. doi: 10.1105/tpc.18.00299

Schilmiller, A. L., Stout, J., Weng, J.-K., Humphreys, J., Ruegger, M. O., & Chapple, C. (2009). Mutations in the cinnamate 4-hydroxylase gene impact metabolism, growth and development in Arabidopsis. In The Plant Journal (Vol. 60, Issue 5, pp. 771–782). Wiley. 10.1111/j.1365-313x.2009.03996.x

Schneider, M., Shrestha, A., Ballvora, A., & Léon, J. (2022). High-throughput estimation of allele frequencies using combined pooled-population sequencing and haplotype-based data processing. Plant methods, 18(1), 34. 10.1186/s13007-022-00852-8

Scossa, F., Alseekh, S., & Fernie, A. R. (2021). Integrating multi-omics data for crop improvement. Journal of plant physiology, 257, 153352. 10.1016/j.jplph.2020.153352

Seo, M., Koshiba, T. Transport of ABA from the site of biosynthesis to the site of action. J Plant Res 124, 501–507 (2011). 10.1007/s10265-011-0411-4

Shannon, P., Markiel, A., Ozier, O., Baliga, N. S., Wang, J. T., Ramage, D., Amin, N., Schwikowski, B., & Ideker, T. (2003). Cytoscape: A Software Environment for Integrated Models of Biomolecular Interaction Networks. In Genome Research (Vol. 13, Issue 11, pp. 2498–2504). Cold Spring Harbor Laboratory. 10.1101/gr.1239303

Shu, K., & Yang, W. (2017). E3 Ubiquitin Ligases: Ubiquitous Actors in Plant Development and Abiotic Stress Responses. In Plant and Cell Physiology (Vol. 58, Issue 9, pp. 1461–1476). Oxford University Press (OUP). 10.1093/pcp/pcx071

Simeão RM, Valle CB do, Alves GF, Moreira DAL, Silva DR da, Araújo D de F, Ferreira RCU, Barrios SCL, Jank L, Caramalac GR, Naka IM, Calixto S and Carvalho J. de (2012) Melhoramento de Brachiaria ruziziensis tetraploide sexual na Embrapa: métodos e avanços. Embrapa, Campo Grande. Documentos 194: 1–32.

Simeão, R, Silva, A., Valle, C., Resende, M. D., & Medeiros, S. (2016). Genetic evaluation and selection index in tetraploid Brachiaria ruziziensis. In H.-P. Piepho (Ed.), Plant Breeding (Vol. 135, Issue 2, pp. 246–253). Wiley. 10.1111/pbr.12353

Simeão, R. M., Valle, C. B., & Resende, M. D. V. (2016). Unravelling the inheritance,QSTand reproductive phenology attributes of the tetraploid tropical grass Brachiaria ruziziensis(Germain et Evrard). In O. A. Rognli (Ed.), Plant Breeding (Vol. 136, Issue 1, pp. 101–110). Wiley. 10.1111/pbr.12429

Simeão RM, Resende MDV, Alves RS, Pessoa-Filho M, Azevedo ALS, Jones CS, Pereira JF and Machado JC (2021) Genomic Selection in Tropical Forage Grasses: Current Status and Future Applications. Front. Plant Sci. 12:665195. doi: 10.3389/fpls.2021.665195

Simeão-Resende, R. M., Casler, M. D., and Resende, M. D. V. (2014). Genomic selection in forage breeding: accuracy and methods. Crop Sci. 54, 143–156. doi: 10.2135/cropsci2013.05.0353

Soneson C, Love MI, Robinson MD (2015). “Differential analyses for RNA-seq: transcript-level estimates improve gene-level inferences.” F1000Research, 4. doi: 10.12688/f1000research.7563.1.

Song, J., Wang, Z. RNAi-mediated suppression of the phenylalanine ammonia-lyase gene in Salvia miltiorrhiza causes abnormal phenotypes and a reduction in rosmarinic acid biosynthesis. J Plant Res 124, 183–192 (2011). 10.1007/s10265-010-0350-5

Stacklies, W., Redestig, H., Scholz, M., Walther, D., and Selbig, J. (2007). pcaMethods a bioconductor package providing PCA methods for incomplete data. Bioinformatics 23, 1164–1167. doi: 10.1093/bioinformatics/btm069

Stakhova, L.; Stakhov, L.; Ladygin, V. Effects of exogenous folic acid on the yield and amino acid content of the seed of Pisum sativum L. and Hordeum vulgare L. Appl. Biochem. Microbiol. 2000, 36, 85–89.

Steinfath, M., Gärtner, T., Lisec, J. et al. Prediction of hybrid biomass in Arabidopsis thaliana by selected parental SNP and metabolic markers. Theor Appl Genet 120, 239–247 (2010). 10.1007/s00122-009-1191-2

Stevens P.F. (2001) Onwards. Angiosperm phylogeny website. Version 12, July 2012 [and more or less continuously updated since]. Available at: http://www.mobot.org/MOBOT/research/APweb/ (Accessed April, 2022).

Thaikua, S., Ebina, M., Yamanaka, N., Shimoda, K., Suenaga, K., and Kawamoto, Y. (2016). Tightly clustered markers linked to an apospory-related gene region and quantitative trait loci mapping for agronomic traits in Brachiaria hybrids. Grassl. Sci. 62, 69–80. doi: 10.1111/grs.12115

Thakral, V., Yadav, H., Padalkar, G., Kumawat, S., Raturi, G., Kumar, V.,…& Singh, M. (2022). Recent Advances and Applicability of GBS, GWAS, and GS in Polyploid Crops. Genotyping by Sequencing for Crop Improvement, 328-354.

Tong, H., & Nikoloski, Z. (2021). Machine learning approaches for crop improvement: Leveraging phenotypic and genotypic big data. In Journal of Plant Physiology (Vol. 257, p. 153354). Elsevier BV. 10.1016/j.jplph.2020.153354

Tuteja, N. (2003). Plant DNA helicases: the long unwinding road. In Journal of Experimental Botany (Vol. 54, Issue 391, pp. 2201–2214). Oxford University Press (OUP). 10.1093/jxb/erg246

Van Lijsebettens, M., Vanderhaeghen, R., De Block, M., Bauw, G., Villarroel, R., & Van Montagu, M. (1994). An S18 ribosomal protein gene copy at the Arabidopsis PFL locus affects plant development by its specific expression in meristems. The EMBO Journal, 13(14), 3378– 3388.

Varshney, R. K. (2021). The Plant Genome special issue: Advances in genomic selection and application of machine learning in genomic prediction for crop improvement. In The Plant Genome (Vol. 14, Issue 3). Wiley. 10.1002/tpg2.20178

Verdoni, N., Mench, M., Cassagne, C., & Bessoule, J.-J. (2001). Fatty acid composition of tomato leaves as biomarkers of metal-contaminated soils. In Environmental Toxicology and Chemistry (Vol. 20, Issue 2, pp. 382–388). Wiley. 10.1002/etc.5620200220

Vigna, B. B. Z., de Oliveira, F. A., de Toledo-Silva, G., da Silva, C. C., do Valle, C. B., and de Souza, A. P. (2016a). Leaf transcriptome of two highly divergent genotypes of Urochloa humidicola (Poaceae), a tropical polyploid forage grass adapted to acidic soils and temporary flooding areas. BMC Genomics 17:910. doi: 10.1186/s12864-016-3270-5

Vigna, B. B. Z., Santos, J. C. S., Jungmann, L., do Valle, C. B., Mollinari, M., Pastina, M. M., et al. (2016b). Evidence of allopolyploidy in Urochloa humidicola based on cytological analysis and genetic linkage mapping. PLoS One 11: e0153764. doi: 10.1371/journal.pone.0153764

Voorrips RE. MapChart: software for the graphical presentation of linkage maps and QTLs. J Hered. 2002;93(1):77–8.

Voss-Fels, K. P., Cooper, M., & Hayes, B. J. (2019). Accelerating crop genetic gains with genomic selection. TAG. Theoretical and applied genetics. Theoretische und angewandte Genetik, 132(3), 669–686.

Wagner, A., Donaldson, L., Kim, H., Phillips, L., Flint, H., Steward, D., Torr, K., Koch, G., Schmitt, U., & Ralph, J. (2008). Suppression of 4-Coumarate-CoA Ligase in the Coniferous Gymnosperm Pinus radiata. In Plant Physiology (Vol. 149, Issue 1, pp. 370–383). Oxford University Press (OUP). 10.1104/pp.108.125765

Waldmann, P., Pfeiffer, C., & Mészáros, G. (2020). Sparse Convolutional Neural Networks for Genome-Wide Prediction. In Frontiers in Genetics (Vol. 11). Frontiers Media SA. 10.3389/fgene.2020.00025

Walter, A., Liebisch, F., & Hund, A. (2015). Plant phenotyping: from bean weighing to image analysis. Plant methods, 11, 14.

Wang, X., Xu, Y., Hu, Z., & Xu, C. (2018). Genomic selection methods for crop improvement: Current status and prospects. In The Crop Journal (Vol. 6, Issue 4, pp. 330–340). Elsevier BV. 10.1016/j.cj.2018.03.001

Wang, X., Shi, S., Wang, G. et al. Using machine learning to improve the accuracy of genomic prediction of reproduction traits in pigs. J Animal Sci Biotechnol 13, 60 (2022). 10.1186/s40104-022-00708-0

Wang, Y., Sun, G., Zeng, Q. et al. Predicting Growth Traits with Genomic Selection Methods in Zhikong Scallop (Chlamys farreri). Mar Biotechnol 20, 769–779 (2018).

Wang, Z., Chapman, D., Morota, G., & Cheng, H. (2020). A Multiple-Trait Bayesian Variable Selection Regression Method for Integrating Phenotypic Causal Networks in Genome-Wide Association Studies. G3 (Bethesda, Md.), 10(12), 4439–4448. 10.1534/g3.120.401618

Wickham, H., and Chang, W. (2016). Package ‘ggplot2’. Vienna: R Foundation for Statistical Computing. doi: 10.1007/978-3-319-24277-4

Winkel-Shirley B (2001) It takes a garden. How work on diverse plant species has contributed to an understanding of flavonoid metabolism. Plant Physiol 127:1399–1404

Wolc, A., & Dekkers, J. (2022). Application of Bayesian genomic prediction methods to genome-wide association analyses. Genetics, selection, evolution: GSE, 54(1), 31. 10.1186/s12711-022-00724-8

Wolfe, C.J., Kohane, I.S. and Butte, A.J. (2005) Systematic survey reveals general applicability of “guilt-by-association” within gene coexpression networks. BMC Bioinform., 6, 227–227. 10.1186/1471-2105-6-227

Worthington, M., Ebina, M., Yamanaka, N., Heffelfinger, C., Quintero, C., Zapata, Y. P., Perez, J. G., Selvaraj, M., Ishitani, M., Duitama, J., de la Hoz, J. F., Rao, I., Dellaporta, S., Tohme, J., & Arango, J. (2019). Translocation of a parthenogenesis gene candidate to an alternate carrier chromosome in apomictic Brachiaria humidicola. In BMC Genomics (Vol. 20, Issue 1). Springer Science and Business Media LLC. 10.1186/s12864-018-5392-4

Worthington, M., Heffelfinger, C., Bernal, D., Quintero, C., Zapata, Y. P., Perez, J. G., De Vega, J., Miles, J., Dellaporta, S., & Tohme, J. (2016). A Parthenogenesis Gene Candidate and Evidence for Segmental Allopolyploidy in Apomictic Brachiaria decumbens. In Genetics (Vol. 203, Issue 3, pp. 1117–1132). Oxford University Press (OUP). 10.1534/genetics.116.190314

Worthington M., Perez J. G., Mussurova S., Silva-Cordoba A., Castiblanco V., Jones C., et al.. (2021). A new genome allows the identification of genes associated with natural variation in aluminium tolerance in *Brachiaria* grasses. J. Exp. Bot. 72, 302–319. doi: 10.1093/jxb/eraa469

Woodward, A. W. (2005). Auxin: Regulation, Action, and Interaction. In Annals of Botany (Vol. 95, Issue 5, pp. 707–735). Oxford University Press (OUP). 10.1093/aob/mci083

Xu, B., Escamilla-Treviño, L. L., Sathitsuksanoh, N., Shen, Z., Shen, H., Percival Zhang, Y.-H., Dixon, R. A., & Zhao, B. (2011). Silencing of 4-coumarate:coenzyme A ligase in switchgrass leads to reduced lignin content and improved fermentable sugar yields for biofuel production. In New Phytologist (Vol. 192, Issue 3, pp. 611–625). Wiley. 10.1111/j.1469-8137.2011.03830.x

Xu, Y., Wang, X., Ding, X. et al. Genomic selection of agronomic traits in hybrid rice using an NCII population. Rice 11, 32 (2018). 10.1186/s12284-018-0223-4

Xue, S., & Barna, M. (2012). Specialized ribosomes: A new frontier in gene regulation and organismal biology. Nature Reviews. Molecular Cell Biology, 13(6), 355– 369. 10.1038/nrm3359

Yan, Z., Huang, H., Freebern, E., Santos, D. J., Dai, D., Si, J., et al. (2020). Integrating RNA-Seq with GWAS reveals novel insights into the molecular mechanism underpinning ketosis in cattle. BMC Genomics 21:489. doi: 10.1186/s12864-020-06909-z

Yang, Y., Han, L., Yuan, Y., Li, J., Hei, N., & Liang, H. (2014). Gene co-expression network analysis reveals common system-level properties of prognostic genes across cancer types. Nature communications, 5, 3231. 10.1038/ncomms4231

Yang, J., Jiang, H., Yeh, C. T., Yu, J., Jeddeloh, J. A., Nettleton, D., & Schnable, P. S. (2015). Extreme-phenotype genome-wide association study (XP-GWAS): a method for identifying trait-associated variants by sequencing pools of individuals selected from a diversity panel. The Plant journal: for cell and molecular biology, 84(3), 587–596.

Yoon, J., Choi, H., & An, G. (2015). Roles of lignin biosynthesis and regulatory genes in plant development. In Journal of Integrative Plant Biology (Vol. 57, Issue 11, pp. 902–912). Wiley. 10.1111/jipb.12422

Yu, H., Zhang, F., Wang, G., Liu, Y., & Liu, D. (2012). Partial deficiency of isoleucine impairs root development and alters transcript levels of the genes involved in branched-chain amino acid and glucosinolate metabolism in Arabidopsis. In Journal of Experimental Botany (Vol. 64, Issue 2, pp. 599–612). Oxford University Press (OUP). 10.1093/jxb/ers352

Zhang, M., Barg, R., Yin, M., Gueta-Dahan, Y., Leikin-Frenkel, A., Salts, Y., Shabtai, S., & Ben-Hayyim, G. (2005). Modulated fatty acid desaturation via overexpression of two distinct ω-3 desaturases differentially alters tolerance to various abiotic stresses in transgenic tobacco cells and plants. In The Plant Journal (Vol. 44, Issue 3, pp. 361–371). Wiley. 10.1111/j.1365-313x.2005.02536.x

Zhang, Z., Ober, U., Erbe, M., Zhang, H., Gao, N., He, J., Li, J., & Simianer, H. (2014). Improving the accuracy of whole genome prediction for complex traits using the results of genome wide association studies. PloS one, 9(3), e93017. 10.1371/journal.pone.0093017

Zhang, H., Wang, X., Pan, Q., Li, P., Liu, Y., Lu, X., Zhong, W., Li, M., Han, L., Li, J., Wang, P., Li, D., Liu, Y., Li, Q., Yang, F., Zhang, Y. M., Wang, G., & Li, L. (2019). QTG-Seq Accelerates QTL Fine Mapping through QTL Partitioning and Whole-Genome Sequencing of Bulked Segregant Samples. Molecular plant, 12(3), 426–437.

Zingaretti LM, Gezan SA, Ferrão LFV, Osorio LF, Monfort A, Muñoz PR, Whitaker VM, Pérez-Enciso M. Exploring Deep Learning for Complex Trait Genomic Prediction in Polyploid Outcrossing Species. Front Plant Sci. 2020 Feb 6; 11:25. doi: 10.3389/fpls.2020.00025. PMID: 32117371; PMCID: PMC7015897.

Zhou, W., Bellis, E.S., Stubblefield, J., Causey, J., Qualls, J., Walker, K. and Huang, X. (2019) Minor QTLs mining through the combination of GWAS and machine learning feature selection. BioRxiv, 712190. 10.1101/712190

Zou, C., Wang, P., & Xu, Y. (2016). Bulked sample analysis in genetics, genomics and crop improvement. Plant biotechnology journal, 14(10), 1941–1955.

