## Supplementary Figures for "Genome-wide family prediction unveils molecular mechanisms underlying the regulation of agronomic traits in *Urochloa ruziziensis*"

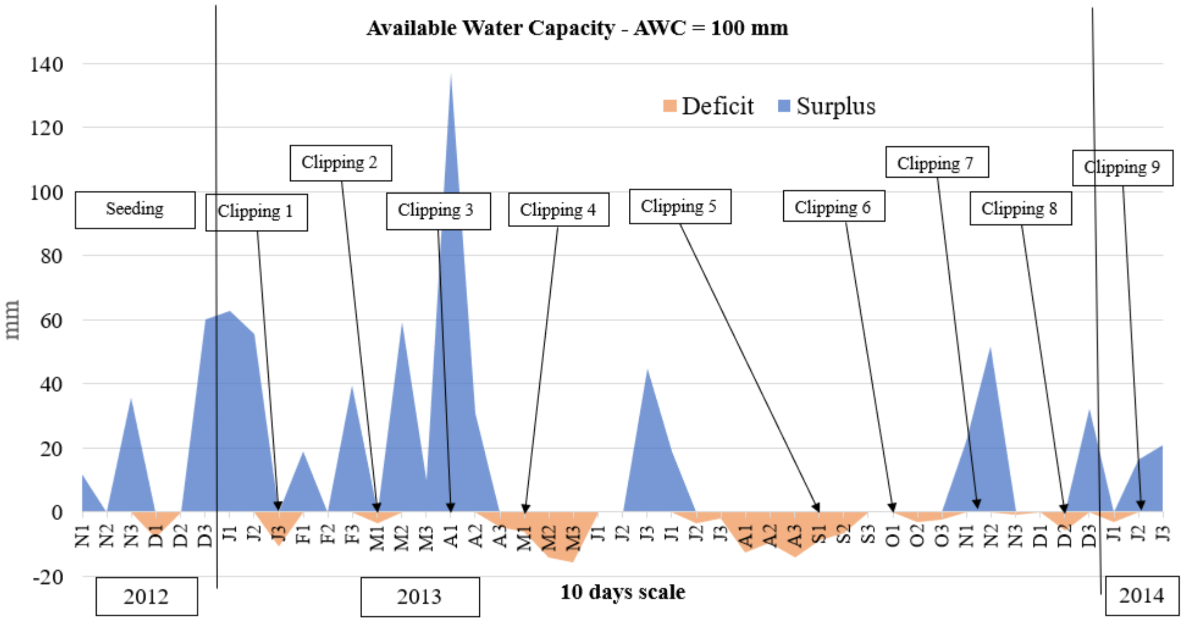


**Supplementary Figure S1.** Climatological water balances using the available water capacity (AWC = 100 mm) on a 10-day scale from November 2012 to January 2014 at the Brazilian Agricultural Research Corporation (Embrapa) Beef Cattle (EBC) station in Campo Grande, Mato Grosso do Sul State, Brazil (20°27’S, 54°37’W, 530 m).

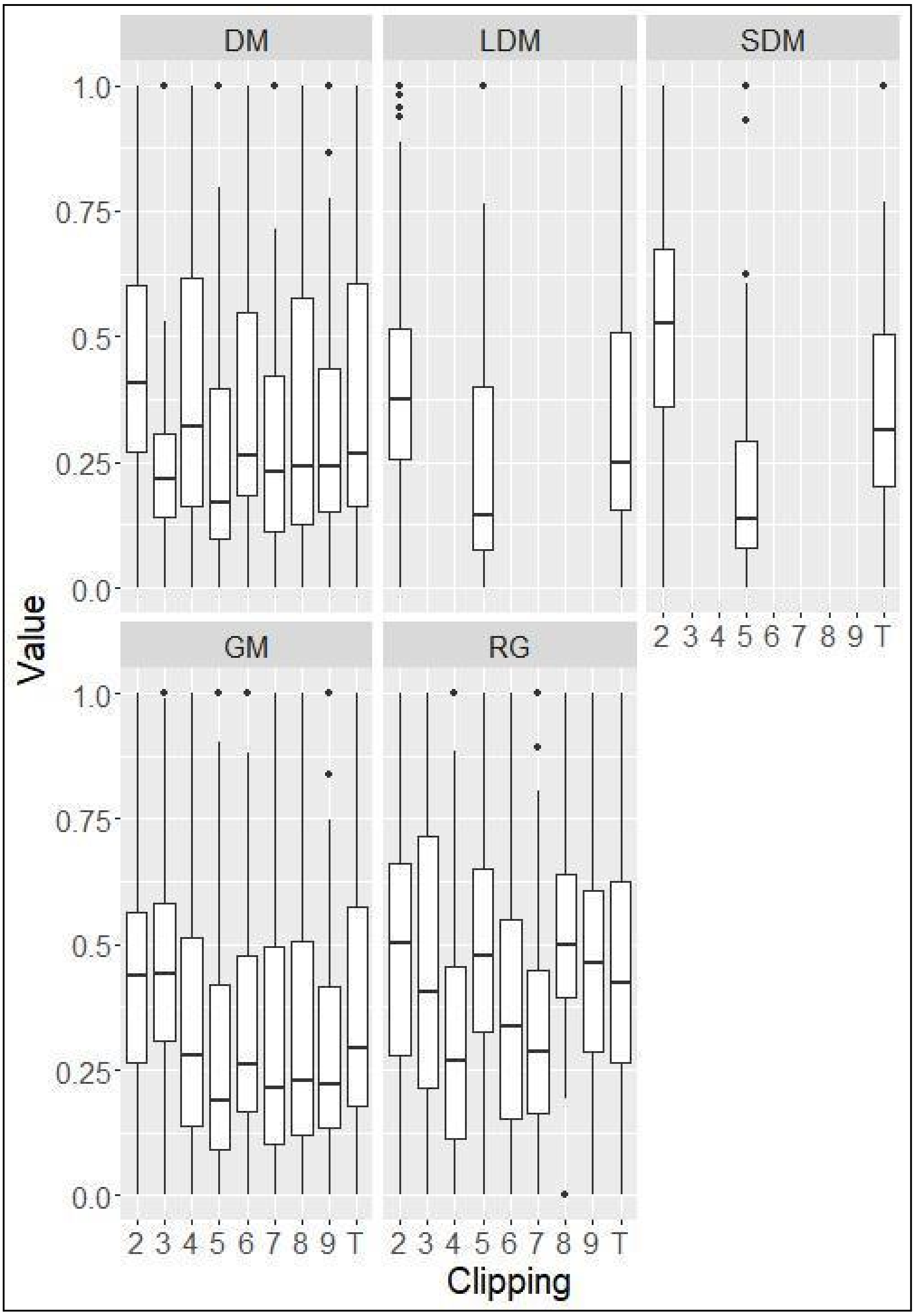


**Supplementary Figure S2.** Box plots of the scaled phenotype clippings. DM, LDM, SDM, GM and RG represent dry matter, leaf dry matter, stem dry matter, green matter and regrowth, respectively.

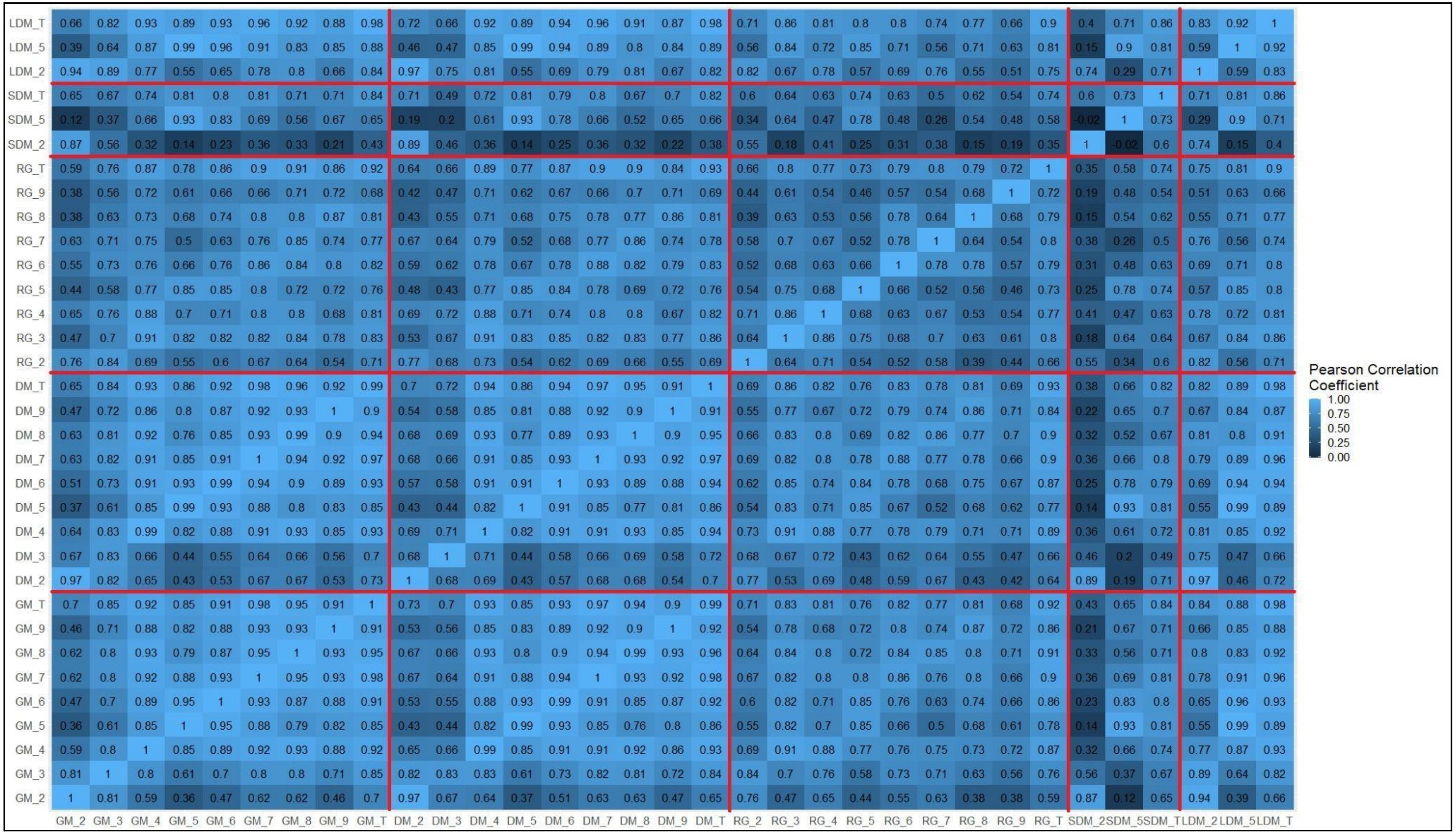


**Supplementary Figure S3.** Heatmap showing the correlation between all trait clippings. Red lines separate different phenotypes. DM, LDM, SDM, GM and RG represent dry matter, leaf dry matter, stem dry matter, green matter and regrowth, respectively.

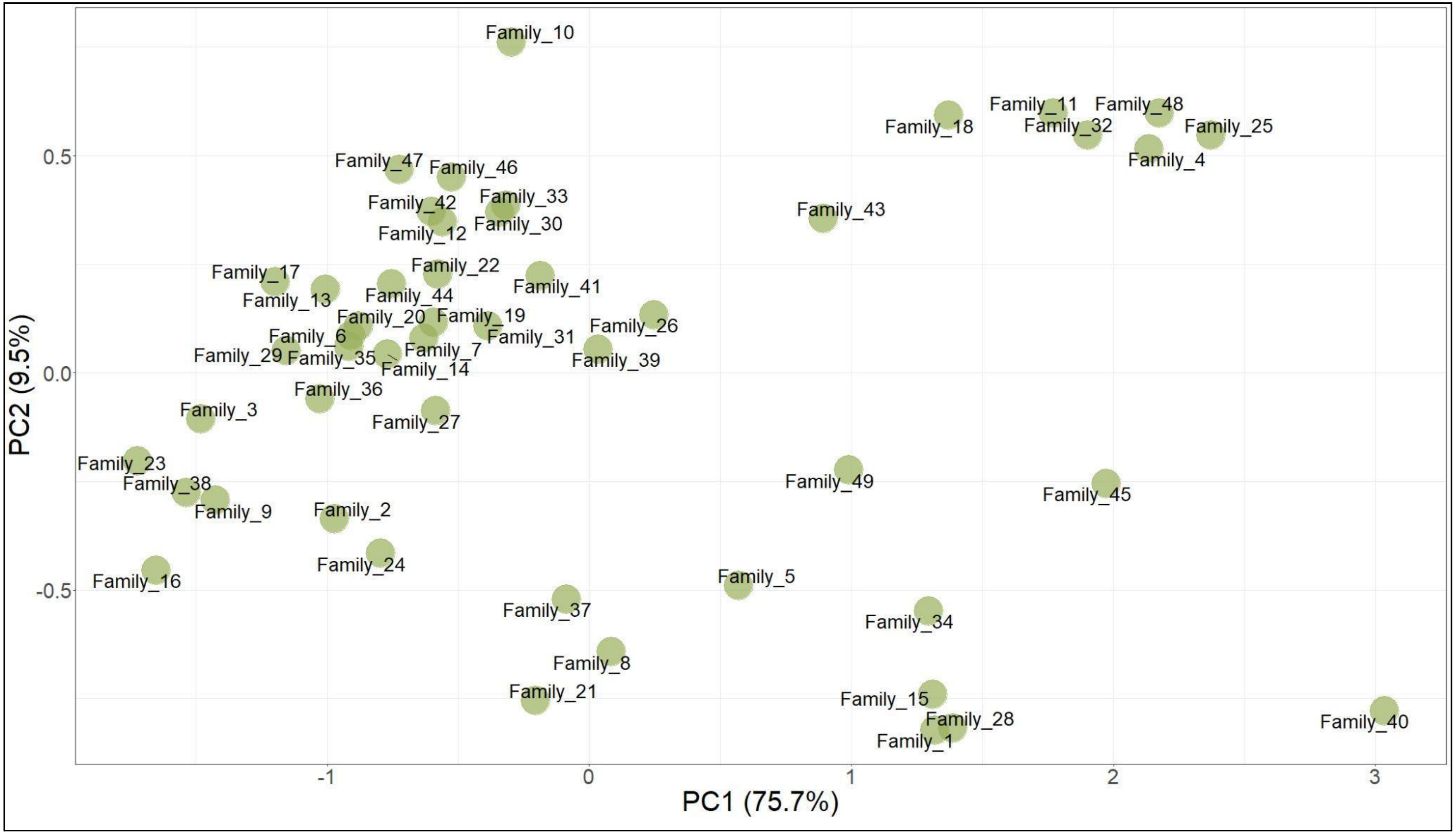


**Supplementary Figure S4.** Principal component analysis scatter plot of the family phenotypes. The axes represent the first and second principal components, which explain 75.7% and 9.5% of the variance, respectively.

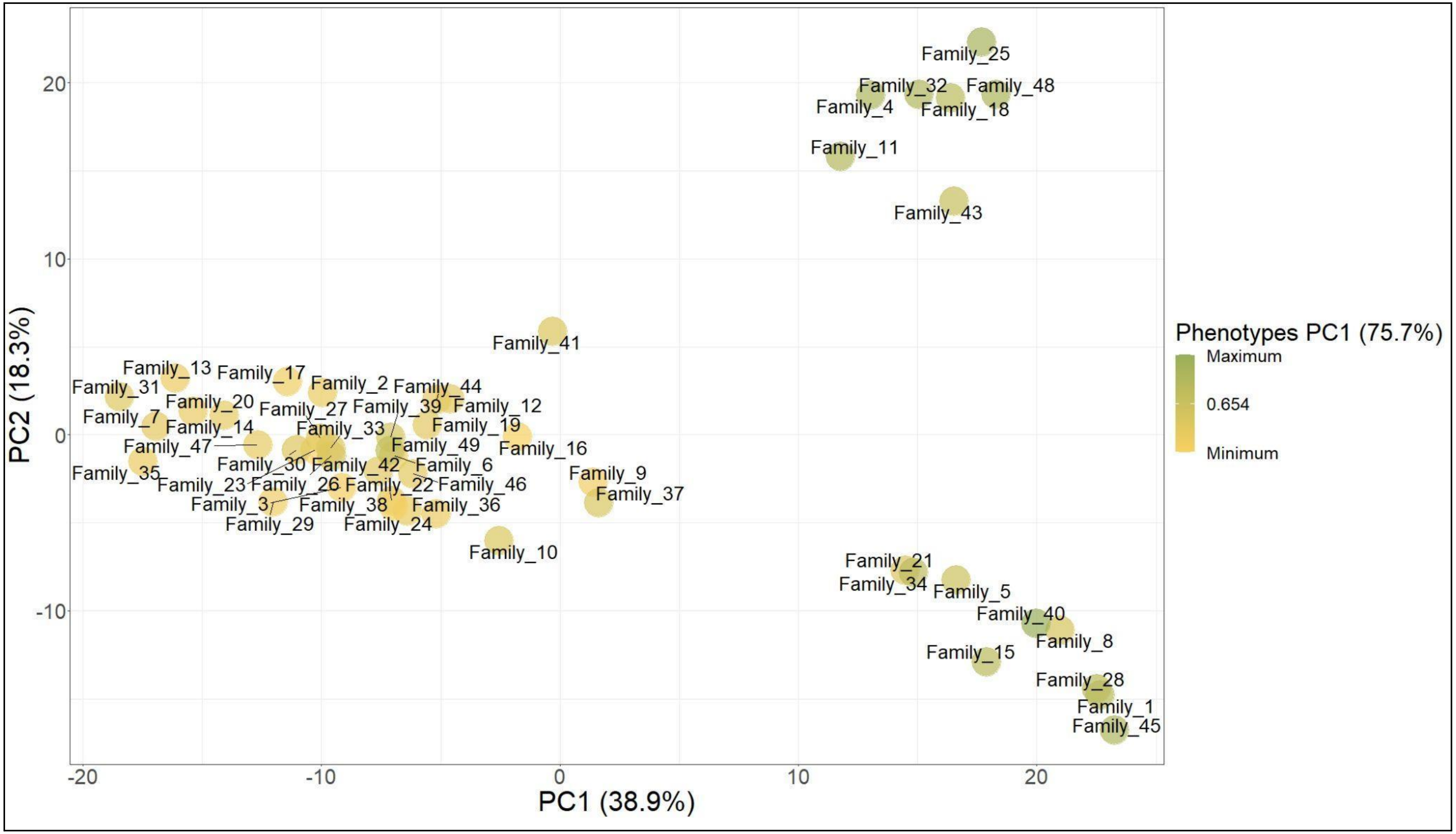


**Supplementary Figure S5.** Principal component analysis scatter plot of family genotyping, with a total of 28,106 markers. The axes represent the first and second principal components, which explain 38.9% and 18.3% of the variance, respectively.

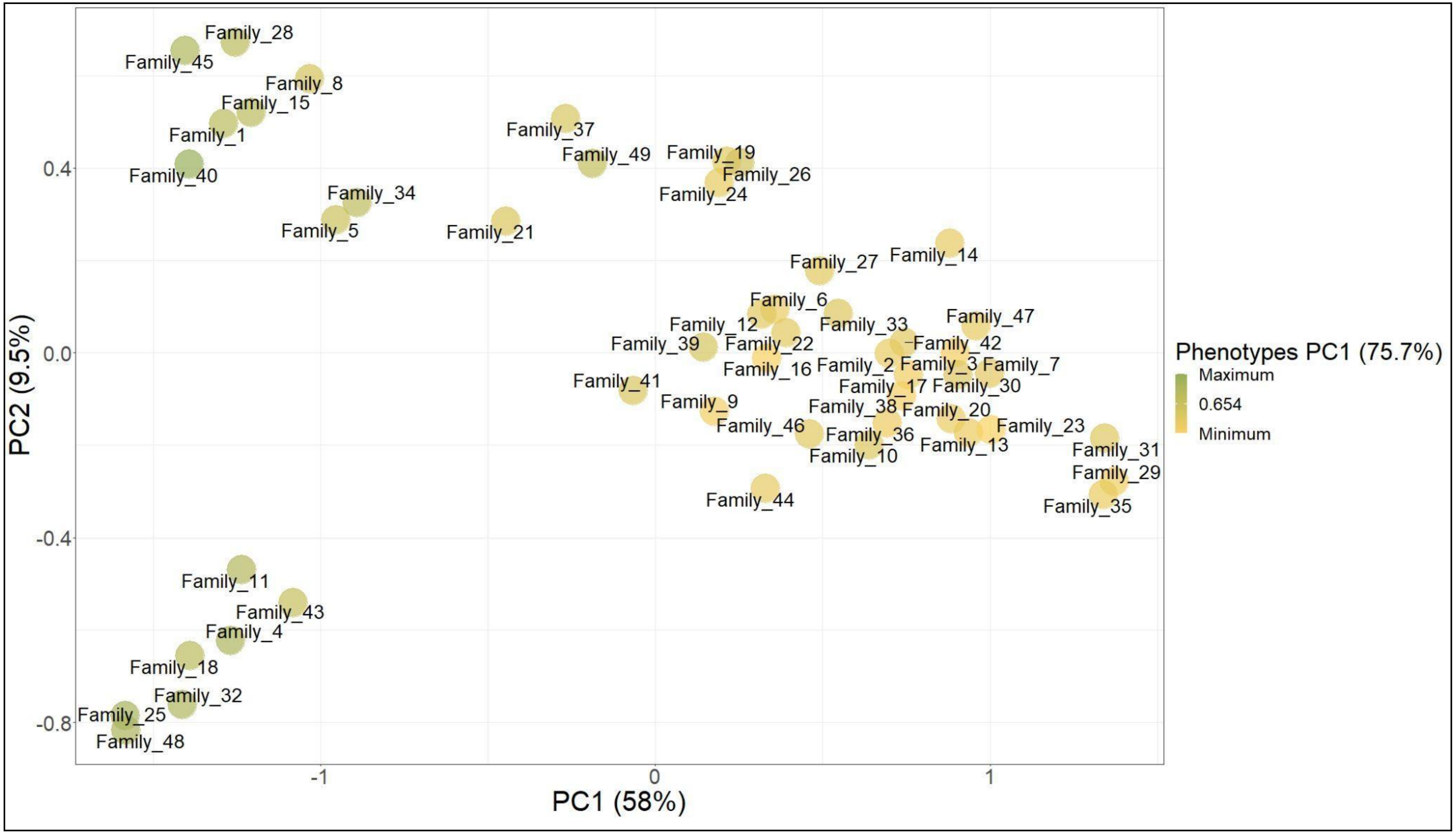


**Supplementary Figure S6.** Principal component analysis scatter plot of family genotyping, considering only the major importance markers (69 markers). The axes represent the first and second principal components, which explain 58% and 9.5% of the variance, respectively.

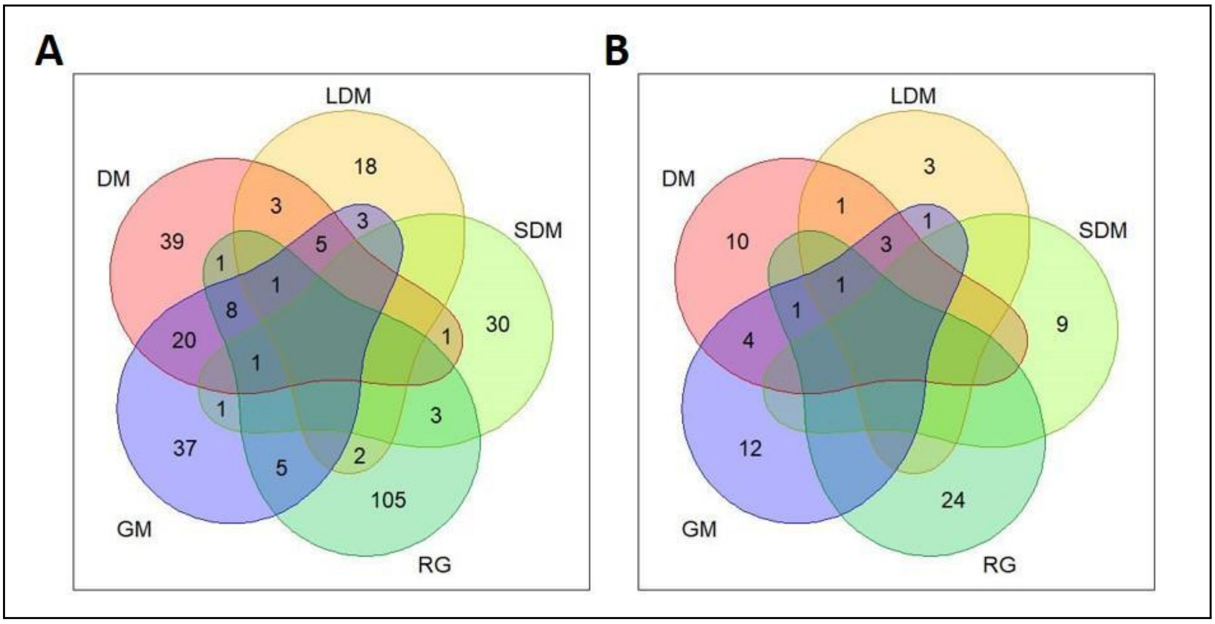


**Supplementary Figure S7.** Venn diagrams showing the logical relationship among the sets of markers identified for each phenotype. (A) A total of 283 markers from the FI-2 dataset and (B) 69 markers selected by the Gini importance condition. DM, LDM, SDM, GM and RG represent dry matter, leaf dry matter, stem dry matter, green matter and regrowth, respectively.

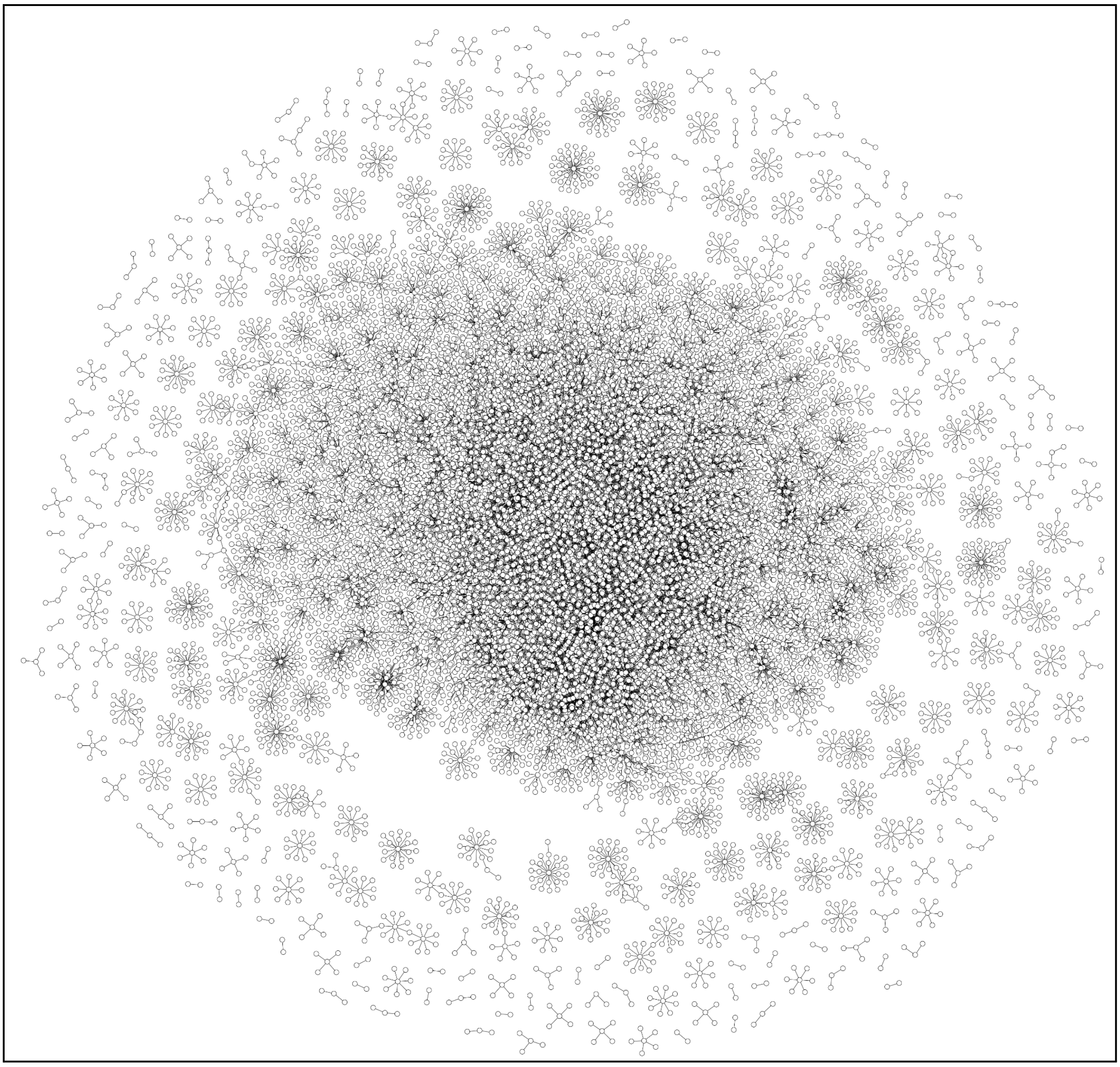

**Supplementary Figure S8.** *Urochloa ruziziensis* gene coexpression network (GCN).
